# Palmitoylated caveolin-1 enables endosome sorting of complex sphingolipids

**DOI:** 10.1101/2025.10.01.679304

**Authors:** SS Schmieder, J Podkalicka, C Viaris de Lesegno, B Han, L Schulz, R Tatituri, M Narendran, L Strieker, J Manzi, AK Kenworthy, P Bassereau, C Lamaze, WI Lencer

**Author notes:** Contributed equally (SS Schmieder, J Podkalicka).

## Abstract

Complex sphingolipids form in the Golgi apparatus and require transport by vesicular carriers to reach the plasma membrane (PM) where they assemble with cholesterol to affect membrane function. The caveolin proteins have been implicated in sphingolipid trafficking but by mechanisms unknown. Here, we found that cells lacking caveolin-1 (Cav1) distributed the complex sphingolipids to the lysosome rather than to the PM. This was not seen in Cavin-1 KO cells, implicating a function for Cav1 independent of caveolae. The defect in trafficking localized to the sorting endosome where the complex sphingolipids failed to enter recycling tubules serving the PM. Sphingolipid trafficking was rescued by over-expression of Cav1, but not by a Cav1 mutant that lacked the S-palmitoylation sites. Thus, noncaveolar and palmitoylated Cav1 appears to act as a chaperone, or selectivity filter, enabling entry of the complex sphingolipids into endocytic recycling tubules to shape the composition of the PM.

## Introduction

Complex sphingolipids comprise up to 20% of total lipids forming the plasma membrane and crucially affect membrane function as structural and signaling components. They encompass sphingomyelin and different glycosphingolipid species that are synthesized from ceramide in the late Golgi compartments where glycosyltransferases or sphingomyelin synthase attach single or multiple sugars or choline to the C1-carbon of ceramide (Hannun and Obeid, 2018). These large hydrophilic headgroups anchor the ceramide domain within the luminal (or exofacial) membrane leaflet, where they are unavailable for interaction with lipid transport proteins in the cytosol (Svistunov et al., 2024). To reach the plasma membrane and other subcellular destinations, the complex sphingolipids depend on vesicular trafficking (Sokoya et al.; Svistunov et al.; van Meer et al., 2008; Young et al., 1992). The cellular mechanisms that govern the vesicular sorting and trafficking of these lipids to their sites of action, however, remain poorly understood.

The complex sphingolipids exhibit a strong affinity for membrane cholesterol to form small, highly dynamic membrane nanodomains (Chiantia and London, 2013; Lingwood and Simons, 2010; Simons and Ikonen, 1997; van Meer et al., 2008). Nanodomains populate the plasma membrane (PM) and the limiting membranes of sub-cellular organelles throughout the exocytic and endocytic networks (Klemm et al.; Simons and Ikonen, 1997; van Meer and Simons, 1982). The strong affinity for cholesterol originates largely from hydrophobic interactions and hydrogen bonding between the ceramide’s amide and the hydroxy-group of cholesterol (Boggs, 1987; Peter Slotte, 2013; Ramstedt and Slotte, 1999; Slotte, 2016), and it is critically affected by the structure of the acyl chain of the sphingolipid (Arumugam et al., 2021) (Schmieder unpublished). We recently discovered a motif within the acyl chain of the glycosphingolipid GM1 that enabled stronger interaction with cholesterol for self-assembly in plasma membrane nanodomains; which we termed the C14* motif (Arumugam et al., 2021; Chinnapen et al., 2012; Schmieder et al., 2022) (Schmieder unpublished). The motif is defined by 14 fully saturated hydrocarbons from the amide bond (Schmieder et al., 2022) (Schmieder unpublished); and it determines the subcellular localization of GM1 by affecting its trafficking within the endocytic system and the plasma membrane (Arumugam et al., 2021; Chinnapen et al., 2012; Schmieder et al., 2022). GM1 with ceramide structures containing the C14*-motif enables strong association with membrane nanodomains and trafficked from the sorting endosome into predominantly the late-endosome/lysosome pathway. In contrast, GM1 species containing ceramide structures without the C14*-motif entered into endosome sorting tubules targeted for recycling to the PM or for retrograde trafficking to Golgi and endoplasmic reticulum (Chinnapen et al., 2012; Schmieder et al., 2022).

Biophysical principles related to lipid shape, membrane stiffness, and phase separation affect the behaviors of the complex sphingolipids in cell membranes and their sub-cellular distributions (Mukherjee and Maxfield, 2000; Roux et al., 2005), but evidence for a dependence in these processes on membrane proteins interacting with the sphingolipids has also been found (Safouane et al., 2010; Sorre et al., 2009; Tian and Baumgart, 2009). The family of caveolin membrane proteins that has long been associated with remodeling and shaping PM organization and composition to affect cell function through association with cholesterol, the complex sphingolipids, and membrane nanodomains (Lamaze et al., in revision; Sinha et al., 2011). In large part, this function originates from the higher order assembly of caveolin with Cavin-1 proteins required for the formation of caveolae – cup-shaped PM invaginations enriched in complex sphingolipids and cholesterol (Hubert et al., 2020b; Kenworthy et al., 2023; Lamaze et al., 2017; Ortegren et al., 2004; Parton et al., 2020; Zhou et al., 2021). The biology of the complex sphingolipids, cholesterol, and caveolae are interdependent. Sphingolipids, for example, regulate caveolae formation and their dynamics (Hubert et al., 2020a; Hubert et al., 2020b; Sharma et al., 2004; Singh et al., 2010). And at the PM, caveolae regulate the membrane dynamics and organization of the sphingolipids (Shvets et al., 2015) and cholesterol (Cheng et al., 2006; Pol et al., 2001), including for example the internalization of glycosphingolipid-binding toxins and viruses leading to disease (Puri et al., 2001; Sotgia et al., 2002; Tagawa et al., 2005; Xing et al., 2020). Evolutionarily however, caveolins predate the formation of caveolae (Hansen and Nichols, 2010) and are known to reside and function independently of caveolae in the PM and in other subcellular compartments, including the Golgi and recycling endosome (Bush et al., 2006; Dupree et al., 1993; Gagescu et al., 2000; Head and Insel, 2007; Khater et al.; Kurzchalia et al., 1992; Lapierre et al., 2012; Lolo et al., 2023; Luetterforst et al., 1999; Pol et al., 1999; Pol et al., 2020; Zheng et al., 2011) – but by mechanisms unknown.

Here, we examine the role of caveolin-1 in trafficking of the complex sphingolipids from Golgi complex to PM and within the sorting endosome after endocytosis. We found a novel function for caveolin-1 acting independently of caveolae in a chaperone-like function for the complex sphingolipids that enabled the entry of these lipids into the narrow and highly curved sorting tubules that emerge from these organelles – thus driving the composition of sphingolipids in the PM at steady state.

## RESULTS

### Cells lacking caveolin-1 have plasma membranes depleted of the complex sphingolipids

To study the role of caveolin-1 (Cav1) in trafficking and sorting of the complex sphingolipids within the secretory and endosomal system, we used two complementary cell-based systems: mouse lung endothelial cells (MLEC) from mice lacking caveolin-1 (Sinha et al., 2011) and the genetically tracktable human skin carcinoma epithelial A431 cells where Cav1 was deleted by CRISPR-Cas9 gene editing (Suppl. Fig. 1A and B). Both cell types normally express high levels of both Cav1 and Cavin-1, and contain caveolae (Sinha et al., 2011; Thomsen et al., 2002).

We first quantified the content of the complex sphingolipid sphingomyelin (SM) at the plasma membrane, which typifies its subcellular distribution at steady state. This was measured in MLEC cells using fluorescently labeled Equinatoxin (EqT) that binds the choline head-group of SM (Suppl. Fig. 1C) (Deng et al., 2016). We observed a striking reduction in plasma membrane (PM) staining for SM in MLEC cells lacking caveolin-1 (MLEC Cav1 KO cells) as compared to MLEC cells derived from WT mice (Fig. 1A and B). The same results were obtained using a different SM binding toxin, OstreolysinA (OlyA) (Suppl. Fig. 1D). Plasma membrane SM levels were recovered in the Cav1 KO cells when transfected with Cav1-RFP. Cells transfected with RFP alone showed no such recovery (Fig. 1C), implicating specificity for Cav1 in rescuing the phenotype.

**Figure 1.**
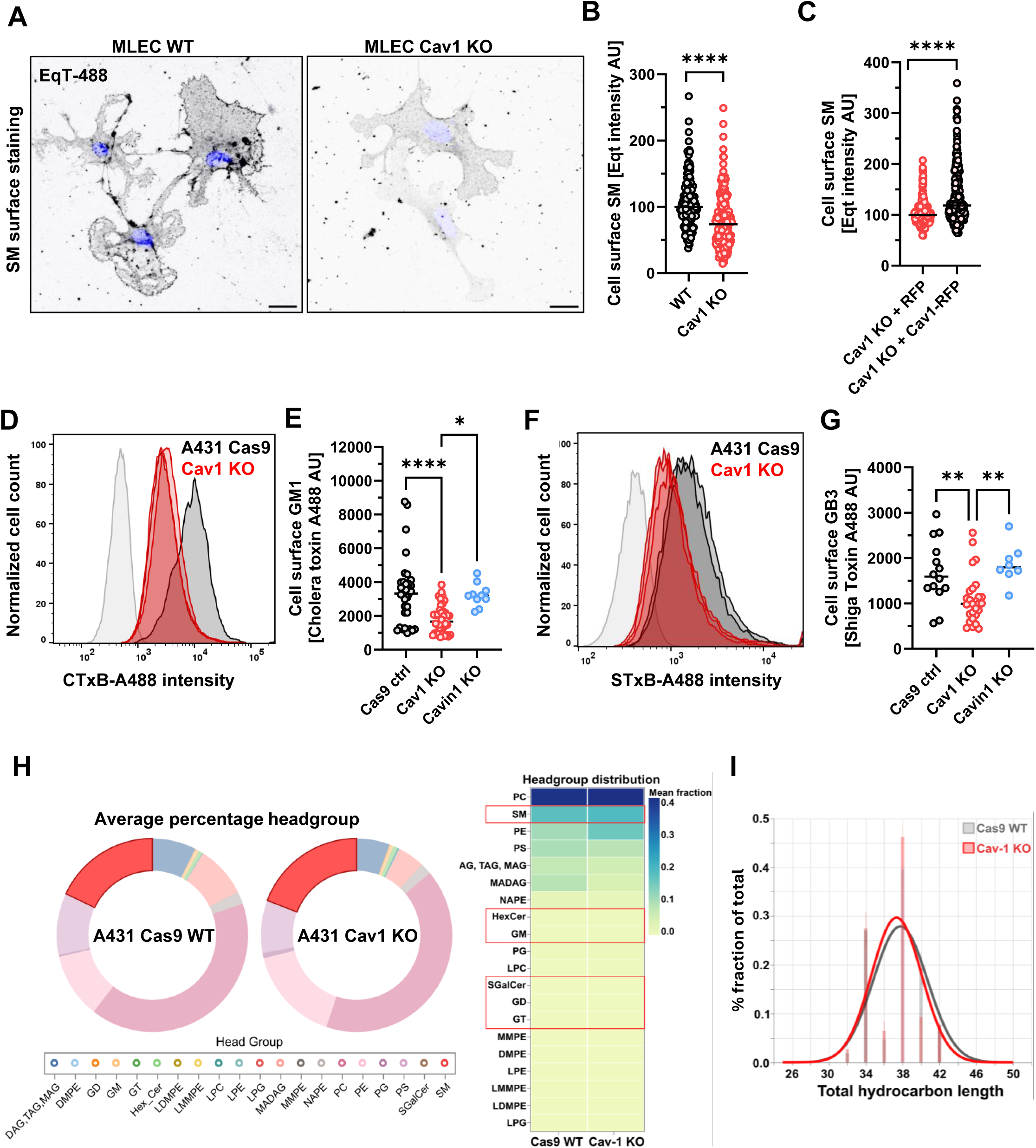
Cell surface depletion of complex sphingolipids in the absence of Caveolin-1. **(A)** Representative cell surface SM staining with EqT-A488 and DAPI (blue) in MLEC WT and Cav1 KO. Scale bar = 20 µm. **(B)** Quantification of imaging in (A), data normalized for WT to 100. Mean displayed. N =3 biological replicates. **(C)** Cell surface SM staining with EqT-A488 in MLEC Cav1 KO cells overexpressing either Cav1-RFP or RFP alone. Data normalized for WT to 100. Mean displayed. N = 3 biological replicates. **(D)** FACS analysis of cell surface GM1 stained with CTxB-A488 in A431 Cas9 WT (grey) and Cav1 KO (red). **(E)** Quantification of (D), dot represents 20.000 cells, mean and SEM, n => 5 biological replicates. **(F)** FACS analysis of cell surface GB3 stained with STxB-A488 in A431 Cas9 WT (grey) and Cav1 KO (red). **(G)** Quantification of (F), dot represents 20.000 cells, mean, n = > 5 biological replicates. **(H)** Lipidomics analysis of A431 Cas9 WT or Cav1 KO cells, n = 3 biological replicates. Doughnut plot shows percentage of lipid species by headgroup in A431. SM in bright red. Heatmap of mean fraction of lipid species by headgroup showing no difference in overall amount of SM, nor the detected GSLs HexCer, GM1, SGalCer, GD or GT. Respective species are encircled by red box. Statistically different lipid species: PE, PS, and MADAG, MAG, DAG and TAG. **(I)** Chain length distribution profile of SM in A431 Cas9 WT (grey) or Cav1 KO (red) shows slight reduction in hydrocarbon length for the Cav1 KO. For all experiments: n.s. p > 0.05, *p < 0.05, **p < 0.01 ***p < 0.001 and ****p < 0.0001.

To test whether Cav1 also affects the abundance of other complex sphingolipids at the PM, we measured the glycosphingolipids GM1 and GB3 in A431 Cav1 KO epithelial cells. We found the same loss of complex sphingolipid content at the PM. In this case, GM1 and GB3 levels were measured quantitatively by FACS using fluorophore-labeled CTxB or STxB to bind GM1 and GB3 respectively (Fig. 1D-G). For both GM1 and GB3, over-expression of Cav1-RFP rescued the depletion of these lipids at the cell surface, consistent with the results we obtained for SM in MLEC cells (Suppl. Fig. 1E and F) and implicating specificity for the loss of Cav1 as cause for the phenotype. A431 cells lacking Cav1, however, also have reduced expression of Cavin-1 (Suppl. Fig. 1G) that is required to stabilize the Cav1 protein coat underlying caveolae structure (Han et al., 2016; Hill et al., 2008; Lamaze et al., 2017; Liu et al., 2022; Liu et al., 2008; Lundmark et al., 2024; Matthaeus and Taraska, 2021). As such, to test if caveolae were required, we generated a Cavin-1 KO in A431 epithelial cells using CRISPR-Cas9 (Suppl. Fig. 1B and G). This cell line continued to express Cav1, though at reduced levels (Supp Fig 1b). In contrast to our results in Cav1 KO cells, we found no significant change in plasma membrane levels for either GM1 and GB3 in A431 cells lacking Cavin-1 (Fig. 1D-G). The result indicates that Cav1 acts independently of Cavin-1 (and thus independently of caveolae) to affect PM levels of the complex sphingolipids.

To determine if the loss of cell surface sphingolipids emerged from a defect in overall sphingolipid biosynthesis, we performed whole cell lipidomics on A431 Cav1 KO and A431 Cas9 WT cells. Both cell lines exhibited comparable overall lipidomes with unaltered mole fractions of the complex sphingolipids (SM, GM1, HexCer, and others) contrary to previous reports from Cav1 KO MEFs (Fig. 1H) (Ariotti et al., 2014; Fernandez-Rojo et al., 2013). Although in the case of SM (the most abundant of the complex sphingolipid so more robustly measured), we observed a small but significant reduction in chain length from C40 to C38, and a small but signifcant increase in the degree of unsaturation when Cav1 was deleted (Fig. 1I and Suppl. Fig. 1H). For the other membrane lipid classes, we detected only minor changes (Fig. 1H and Suppl. Fig. 1I and J). Closely similar results were obtained from the lipidomic profiling of MLEC WT and Cav1 KO cells (data to be uploaded to Lipid Maps). The A431 Cav1 KO cells also displayed elevated levels of cholesterol esters compared to the A431 Cas9 WT and Cavin-1 KO cell lines (Suppl. Fig. 1L). But we did not detect any change in PM cholesterol levels as assessed by filipin staining (Suppl. Fig. 1M). We also analyzed protein levels for key biosynthetic enzymes in the de novo synthesis of the sphingolipids, including serine palmitoyltransferase 2 (SPTLC2), ceramide synthase 5 (CerS5), ceramide synthase 2 (CerS2), ceramide glucosyltransferase (UGCG) and beta-1,4-galactosyltransferase 6 (B4GALT6). Expression of these enzymes was unaltered by deletion of Cav1 or Cavin-1 in A431 cells (Suppl. Fig. 1K).

Thus, the total cellular content of the complex sphingolipids remained unchanged in the caveolin-1 KO cell lines; and a general reduction in total cellular sphingolipids cannot explain the depletion of the sphingolipids at the cell surface observed in either A431 or MLEC Cav1 KO cells. These results implicate a defect in membrane trafficking as a cause for the loss of the complex sphingolipids at the PM in cells lacking Cav1.

### Cells lacking caveolin-1 exhibit an altered post-Golgi secretory pathway for SM

We next asked whether the decreased levels of cell surface SL resulted from defects in biosynthetic trafficking. To test this, we quantitatively measured SM trafficking from its site of production in the Golgi to the plasma membrane using EqT applied to the retention-<u>u</u>sing-<u>s</u>elective-<u>h</u>ook (RUSH) approach (Boncompain et al., 2012). To synchronize EqT exit from the ER in WT and Cav1 KO MLEC cells, we exogenously expressed a modified EqT-GFP containing a streptavidin-binding peptide (EqT-SBP) that binds an exogenously expressed streptavidin-KDEL anchor protein localized primarily to the lumen of the ER. Release of EqT from the streptavidin-KDEL anchor in the ER into the secretory pathway was induced by the addition of biotin (Fig. 2A) (Boncompain et al., 2012).

**Figure 2.**
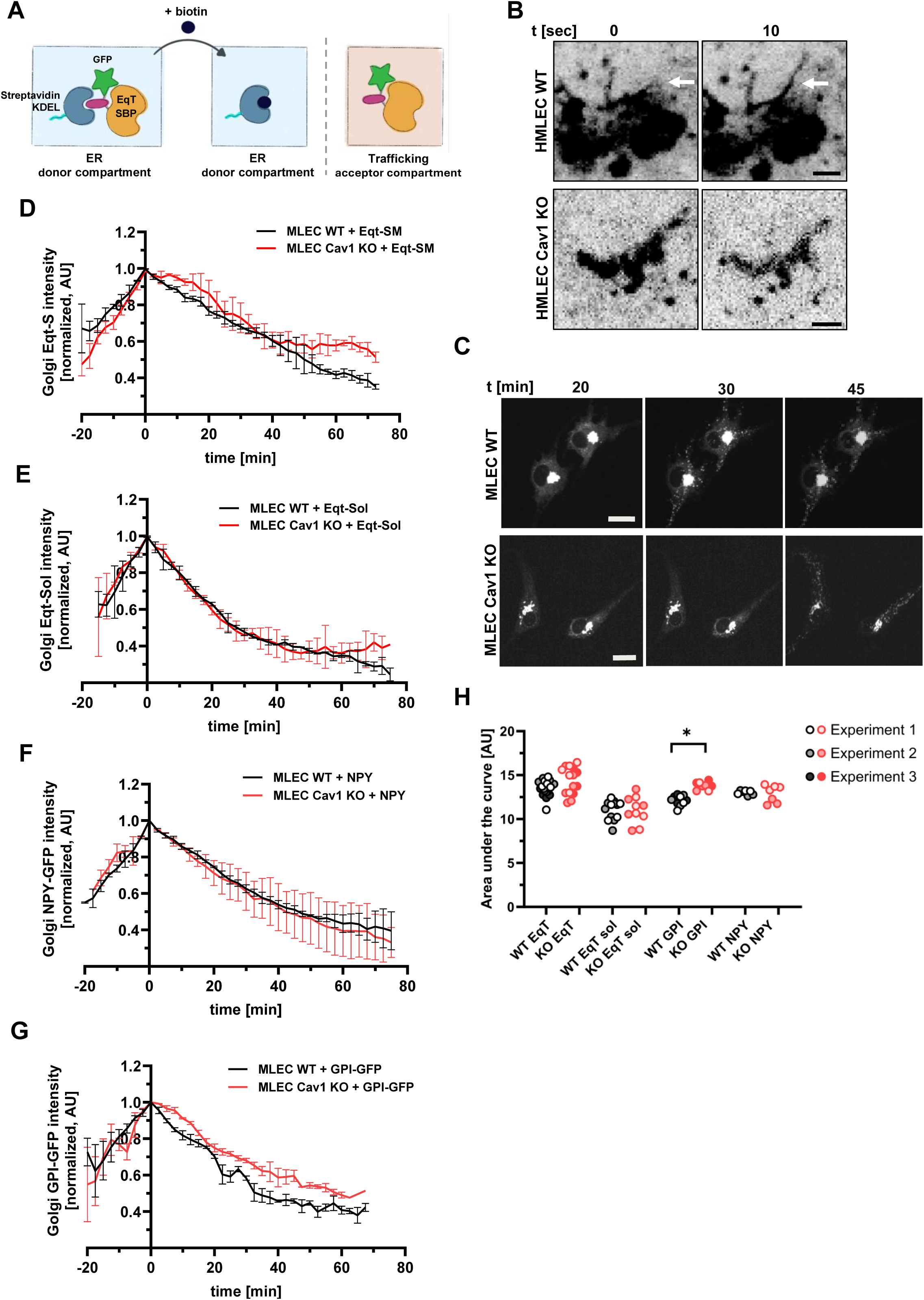
EqT-Rush time course shows moderate reduction in Golgi exit for EqT-SM complex and GPI anchored protein in MLEC Cav1 KO. **(A)** Cartoon depicting RUSH experimental approach. **(B)** MLEC WT displays frequent EqT-SM Golgi tubules which are absent in the Cav1 KO. RUSH image of EqT-GFP. Scale bar = 1 µm. **(C)** Post-Golgi time course for secretory EqT-RUSH trafficking in MLEC WT and Cav1 KO cells. Scale bar = 20 µm. **(D)** Kinetics and quantification of EqT-RUSH trafficking out of the Golgi and into the secretory pathway in MLEC WT (black lines) and Cav1 KO cells (red lines) for N = 3 biological replicates. Time point zero is defined as maximal EqT signal intensity co-localizing to Golgi marker αManII. **(E)** Quantification of non-SM binding EqT-sol-RUSH trafficking MLEC WT (black lines) and Cav1 KO cells (red lines) for N = 2 biological replicates. Experiments defined as in (D). **(F)** Quantification of soluble, secreted protein NPY-GFP-RUSH trafficking MLEC WT (black lines) and Cav1 KO cells (red lines) for N = 2 biological replicates. Experiments defined as in (D). **(G)** Quantification of GPI-GFP-RUSH trafficking MLEC WT (black lines) and Cav1 KO cells (red lines) for N = 3 biological replicates. Experiments defined as in (D). **(H)** Area under the curve (AUC) measurements of RUSH time courses for timepoints 0 to 75min for EqT in (D), EqT-sol in (E), secretory protein NPY in (F) and GPI-GFP in (G). For all experiments: n.s. p > 0.05, *p < 0.05, **p < 0.01 ***p < 0.001 and ****p < 0.0001.

Trafficking into and out of the Golgi was quantified by imaging in live cells over time, with the Golgi compartment defined by stable expression of the cis-Golgi protein α-mannosidase II (MAN II) tagged with mCherry (Suppl. Fig. 2A). The rates of exit from the Golgi for all studies were normalized to the maximal EqT-Golgi colocalization signal (typically obtained about 20 min after the addition of biotin). An apparent defect in the EqT Golgi/post-Golgi secretory pathway was observed in MLEC Cav1 KO cells in agreement with previous reports (Carpentier et al., 2025). First, we observed the absence of elongated EqT-SBP secretory tubules emanating from the Golgi – as compared to WT MLEC cells (Fig. 2B). Post-Golgi EqT secretory vesicles in the MLEC Cav1 KO cells were also delayed in exit from the Golgi, fewer in number (Fig. 2C and Suppl. Fig. 2B and C), of larger size (Suppl. Fig. 2D), and they moved more slowly (Suppl. Fig. 2E) over smaller distances (Suppl. Fig. 2F). We could not detect such size defects in the Golgi/post-Golgi secretory pathway when we analyzed post-Golgi vesicle size for the soluble, non-binding SM mutant (EqT-sol, Suppl. Fig. 2G). In spite of these observed defects, however, the absence of caveolin-1 in MLEC Cav1 KO cells did not lead to a statistically significant difference in EqT-SBP delivery to the PM compared to MLEC WT cells (Suppl. Fig. 2H and I), implying that complex sphingolipids traffic normally to the plasma membrane in Cav1 KO cells.

When the rate (kinetics) of SM exit from the Golgi was studied, however, we found a small - though not significant - delay in export of EqT-SBP (Fig. 2D and H). This was not apparent for the soluble mutant equinotoxin that cannot bind SM (EqT_sol_-SBP), or for the soluble secreted protein neuropeptide Y (NPY also labeld with the RUSH tag) (Fig. 2E, F, and H). Subsequent experiments using a RUSH-tagged GPI-anchored GFP to model a different lipid-anchored nanodomain-associated cargo showed a delay in Golgi exit in Cav1 KO cell analogous to what we found for EqT-SM – but in this case the effect achieved statistical significance (Fig. 2G and H). Given these results, we conclude MLEC cells lacking caveolin-1 likely display a defect in the post-Golgi secretory pathway for both SM and GPI-anchored nanodomain cargoes – though the magnitude of the effect is small overall.

### CAV1 KO cells aberrantly redistribute the Sphingolipids into the lysosomal pathway

We next evaluated the subcellular locations of SM at steady state by staining for SM in MLEC WT and Cav1 KO cells using fluorescently-labeled EqT (Fig. 3A). The Cav1 KO MLEC cells, compared to WT cells, exhibited a striking increase in staining for intracellular EqT (Fig. 3A and B). When counter-stained with the lysosomal marker Lamp1, we found a significant increase of EqT co-localizing with Lamp1 positive vesicles in MLEC Cav1 KO cells (Fig. 3C and D). Similar observations of enhanced transport to the lysosome for excess exogenously applied glycosphingolipids were made in mouse embryonic fibroblasts (MEF) knocked out for Cav1 cells (Shvets et al., 2015). In the case of our studies, transient overexpression of Cav1-RFP, but not RFP alone, reduced the extent of co-localization between Lamp1 and EqT (Supp. Fig. 3A). Notably however, in live cells, none of the post-Golgi secretory vesicles containing EqT-SM complexes were observed to fuse directly with lysotracker labeled lysosomes (Fig. 3E and F and Suppl. Video 1) – suggesting that the trafficking of SL from the Golgi complex to the lysosome must be indirect, presumably via endocytic uptake and trafficking to the lysosome after transport to the PM.

**Figure 3.**
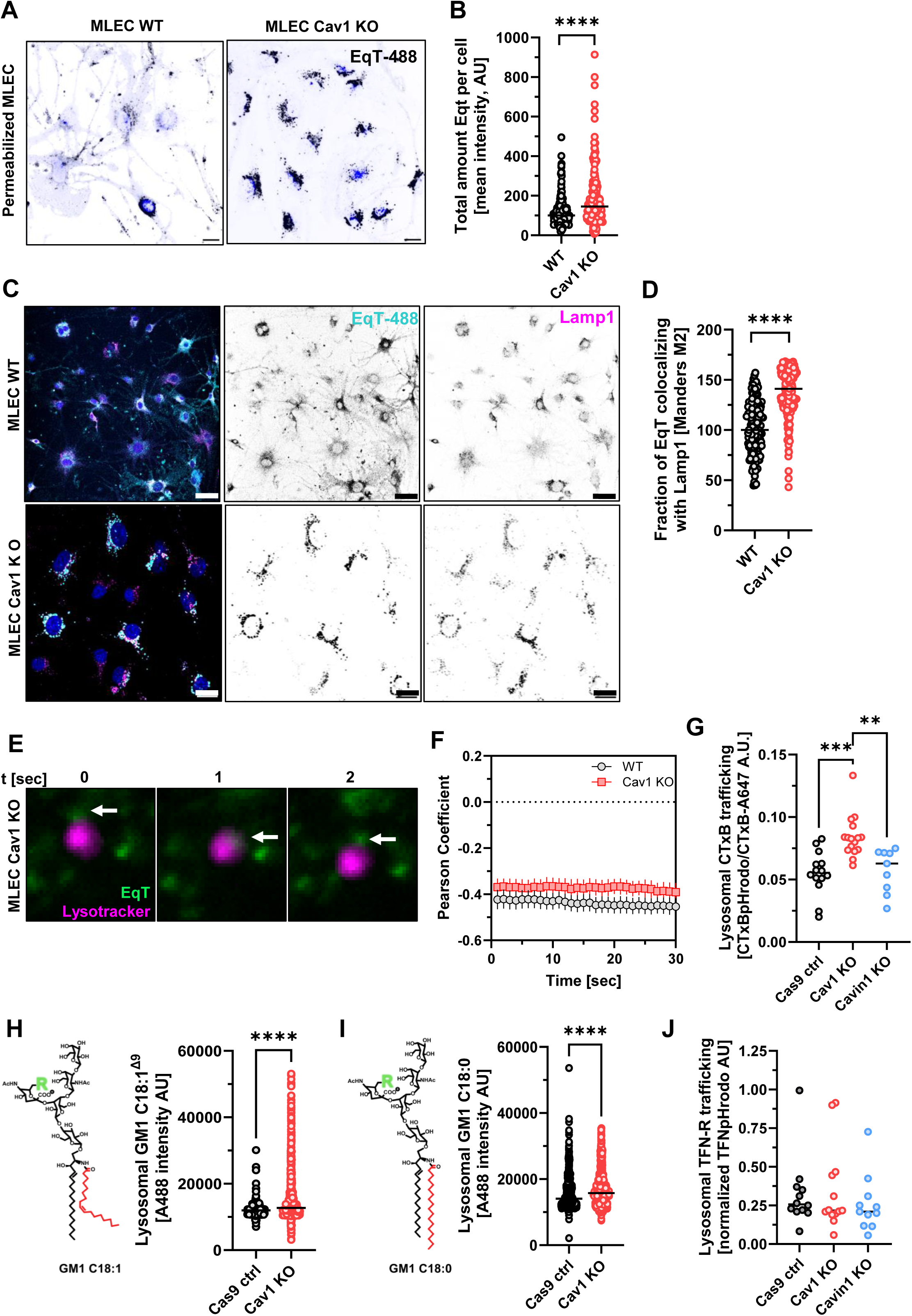
Redistribution of SLs to lysosomes in Cav1 KO cells. **(A)** Permeabilized MLEC WT or Cav1 KO show increase in total intracellular EqT-A488 in Cav1 KO. DAPI staining in blue. Scale bar = 20 µm. **(B)** Quantification of (A) total EqT staining in saponin permeabilized cells for n = 3 biological replicates. **(C)** Immunostaining for SM and lysosomal marker Lamp-1 in MLEC WT or Cav1 KO cells. Scale bar = 20 µm. **(D)** Quantification of (C) for n = 3 replicates. **(E)** EqT containing vesicles (green) do not fuse with lysotracker stained lysosomes (magenta) in MLEC Cav1 KO cells. **(F)** Quantification of (E) by Pearson’s coeffiecient. **(G)** Lysosomal trafficking of CTxB-pHrodo (binding GM1) measured by FACS is increased in the A431 Cav1 KO compared to A431 Cas9 WT and Cavin-1 KO. Lysosomal CTxB-pHrodo fluorescence is normalized to plasma membrane binding of CTxB-A647. N ≥ 7 biological replicates. **(H)** Amount of fluorescently labeled GM1 C18:1^Δ9^ colocalizing to the lysosome is increased in A431 Cav1 KO compared to Cas9 WT. N =>5 biological replicates. **(I)** Amount of fluorescently labeled GM1 C18:0 colocalizing to the lysosome is increased in A431 Cav1 KO compared to Cas9 WT. N =>5 biological replicates. **(J)** Lysosomal trafficking of the recycling cargo TFN-R measured by TFN-pHrodo and FACS is not impaired in A431 Cav1 KO or Cavin-1 KO compared to A431 Cas9 WT. N ≥ 11 biological replicates. For all experiments: n.s. p > 0.05, *p < 0.05, **p < 0.01 ***p < 0.001 and ****p < 0.0001.

To test for enhanced lysosomal transport after endocytosis, we studied trafficking of the glycosphingolipid GM1 in A431 cells. A431 cells lacking Cav1 also displayed increased number of lysosomes, but with slightly reduced size, as measured by lysotracker staining (Suppl. Fig. 3B and C). We quantified transport of endogenous cell surface GM1 into the lysosome by FACS using pH-sensitive pHrodo-labeled CTx-B. A431 Cav1 KO cells exhibited a greater fluorescence signa, implicating enhanced trafficking of CTxB-pHrodo into the acidic late endosome/lysosomal pathway as compared to A431 Cas9 control cells (Fig. 3G). No such amplification in transport of CTxB-GM1 between the PM and lysosome was observed in cells lacking Cavin-1, ruling out a requirement for caveolae in this process. (Fig. 3G).

In the experiments above, we monitored the trafficking of endogenous SLs, which contain a range of fatty acid structures in their ceramide moieties (Hannun and Obeid, 2018; Schmieder et al., 2022). To measure endocytic trafficking of GM1 directly, as well as to test how differences in fatty structure might contribute to the sorting defect implicated in Cav1 KO cells, we used two species of GM1 taken from our ceramide-structural library (Schmieder et al., 2022). In one species, the ceramide carries a fatty acid with double bond at position Δ9 – termed GM1 C18:1 (Fig. 3H cartoon). The cis double bond at position Δ9 disrupts the C14*-motif for nanodomain association and allows trafficking of this GM1 species into all endosomal pathways (recycling, lysosomal, retrograde and transcytotic) (Chinnapen et al., 2012; Garcia-Castillo et al., 2018; Schmieder et al., 2022). The other species, GM1 C18:0, carries a fully saturated acyl chain (Fig. 3I cartoon) that contains the C14*-motif and enables nanodomain association (Arumugam et al., 2021; Schmieder et al., 2022). This GM1 species is strongly sorted into the lysosome pathway (Chinnapen et al., 2012; Schmieder et al., 2022). Both GM1 lipids contained the same oligosaccharide head group fluorescently labeled with Alexa-488.

When trace amounts of these lipids were incorporated into the PM of A431 cells, we found enhanced transport from PM to lysosome for both GM1 species in Cav1 KO cells, implicating a defect in endosomal recycling independent of assembly in nanodomains (Fig. 3H and I). To test whether this effect was specific to SL sorting or due to a general feature of the sorting endosome in Cav1 KO cells, we examined the trafficking of the rapidly recycling transferrin receptor TfnR and the lysosome-targeted low density lipoprotein receptor LDLR (Fig. 3J and Suppl. Fig. 3D). We did not observe amplified transport into the lysosome for either membrane protein. We also observed no effect of Cav1 deletion on endocytosis for the fluid phase solute 4 kDa dextran (Suppl. Fig. 3E). These results are consistent with our observations of SM trafficking in MLEC cells, and they implicate an endosomal sorting defect specific to the complex sphingolipids caused by the loss of Cav1.

### Sorting of GM1 into endosome recycling tubules requires Cav1

The apparent redistribution of exogenously added GM1 from the PM to the lysosome in Cav1 KO cells implies a defect within the early/sorting endosome or recycling pathway. To test for this, we tagged the two GM1 species C18:1 and C18:0 with biotin and quantitatively measured their distribution in the PM of A431 and HMLEC cells over time. Cell surface distribution was measured by FACS using fluorescence-labeled streptavidin to bind the biotin-GM1 located at the cell surface at different time points (Fig. 4A). The level of retention of exogenously applied GM1 content in the PM over time was taken as evidence for endosome recycling of GM1 (as we defined before (Schmieder et al., 2022)).

**Figure 4.**
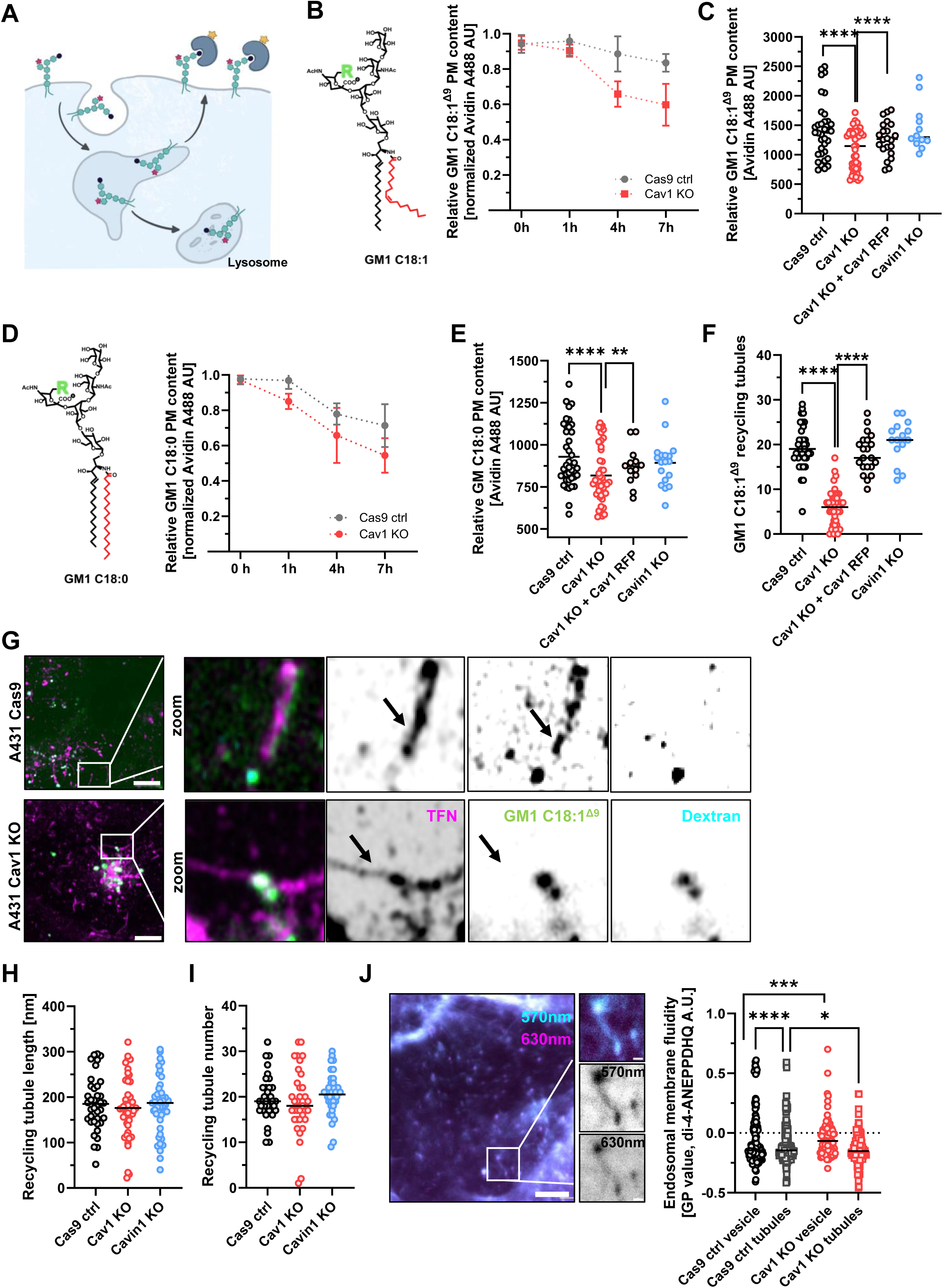
Sphingolipid plasma membrane recycling is impaired in Cav1 KO. **(A)** Cartoon for structural GM1 recycling assay in A431. After endocytosis GM1 is either recycled back to the PM via recycling endosomal tubules or matures with the endosome towards the degradative pathway/lysosome. **(B)** Examplary recycling assay time course in A431 Cas9 WT and Cav1 KO for the functionalized glycosphingolipid GM1 with C18:1^Δ9^ fatty acid (red). Modification of GM1 with either fluorophore or biotin in the glycan headgroup (see GM1 cartoon). A431 Cav1 KO displays reduced recycling to the PM of GM1 C18:1^Δ9^ after endocytosis. **(C)** Quantification of GM1 C18:1^Δ9^ plasma membrane recycling 4 h post loading for in A431 Cas9 WT, Cav1 KO, Cav1 KO reconstituted with Cav1-RFP and Cavin-1 KO. N ≥ 12 biological replicates, dot represents 20.000 cells. **(D)** Glycosphingolipid GM1 with C18:0 fatty acid (red), modified as in (B). Exemplary time course for plasma membrane recycling of GM1 C18:0 in A431 Cas9 WT and Cav1 KO. Cav1 KO displays reduced recycling of GM1 C18:0. **(E)** Quantification of GM1 C18:0 plasma membrane recycling 4 h post loading for in A431 Cas9 WT, Cav1 KO, Cav1 KO reconstituted with Cav1-RFP and Cavin-1 KO. N ≥ 12 biological replicates, dot represents 20.000 cells. **(F)** Quantification of GM1 C18:1^Δ9^ positive TFN recycling tubules for A431 Cas9 WT, Cav1 KO, Cav1 KO reconstituted with Cav1-RFP and Cavin- 1 KO. N ≥ 3 biological replicates. **(G)** GM1 C18:1^Δ9^ colocalizes to TFN (magenta) positive recycling tubules in the A431 Cas9 but is absent from TFN (magenta) positive recycling tubules in A431 Cav1 KO (black arrows). Scale bar = 5 µm. **(H)** Quantification of TFN positive recycling tubule length in A431 Cas9, Cav1 and Cavin-1 KO. N ≥ 3 biological replicates. **(I)** Quantification of TFN positive recycling tubule numbers in A431 Cas9, Cav1 and Cavin-1 KO. N ≥ 3 biological replicates. **(J)** Quantification of membrane order (GP value) measured by the polarity sensitive dye di-4-ANEPPDHQ in TFN positive recycling vesicles and tubules for A431 Cas9 WT and Cav1 KO displays decreased order in A431 Cav1 KO tubules. N ≥ 4 biological replicates. For all experiments: n.s. p > 0.05, *p < 0.05, **p < 0.01 ***p < 0.001 and ****p < 0.0001.

We found that both GM1 species were depleted from the PM more rapidly and to a greater extent in Cav1 KO cells as compared to WT cells loaded with the same lipids respectively (Fig. 4B-E). The amplified depletion of both GM1 species from the PM was rescued by overexpression of Cav1-mCherry (Fig. 4C and E). The same results were obtained for GM1 applied to the MLEC cell lines (Suppl. Fig. 4A and B). Consistent with our observations on the trafficking of SLs in the secretory pathway, A431 cells lacking Cavin-1 phenocopied WT A431 cells with respect to GM1 endosome recycling (Fig. 4C and E). These observations implicate a defect in sorting GM1 into endosome recycling tubules caused by the absence of non-caveolar Cav1. This defect in endosome sorting cannot be explained by alterations in the biology of membrane nanodomains as both the C18:1 and C18:0 GM1 species were affected.

To determine the site of the sorting defect, we visualized endosome sorting of GM1 using the fluorescently-labeled GM1-C18:1 species in combination with fluorescently-labeled Tfn (marking a well-studied receptor-mediated endosome recycling pathway) and dextran (a fluid-phase uptake and late-endosome/lysosome pathway marker). Vesicles containing all three probes functionally defined the early sorting endosome compartment (Fig. 4F and G). Sorting tubules emanating from the early sorting endosome and containing both Tfn-R and GM1-C18:1 were readily visualized in WT A431 cells (Fig. 4F and G) and in cells lacking Cavin-1 (Fig. 4F). In contrast, a striking absence of GM1-C18:1 in Tfn-R positive recycling tubules was observed in the A431 Cav1 KO compared to the A431 WT cells (Fig. 4F and G). Sorting of GM1-C18:1 into recycling endosomal tubules of A431 Cav1 KO cells was restored by overexpressing Cav1-mCherry (Fig. 4F). The depletion of GM1 C18:1 from recycling endosomal tubules in Cav1 KO cells implicates a defect in SL sorting by the recycling endosome - consistent with our earlier results where we observed a depletion of the GM1 species from the plasma membrane (Fig. 4B-E). We did not, however, detect any morphological differences to the recycling endosomal compartment overall (Fig. 4H and I), unlike what we found in the post-Golgi secretory pathway in MLEC cells lacking Cav1 (Suppl. Fig. 2B-F).

The finding that complex sphingolipids (presumably in complex with cholesterol) failed to enter endosome recycling tubules in Cav1 KO cells predicts that these sorting tubules should display greater membrane fluidity compared to endosome sorting tubules in control WT cells (Rawicz et al., 2008). Likewise, the retention of the complex sphingolipids in the sorting endosome itself (en route to the lysosome) predicts that these vesicles should display less membrane fluidity. To test these predictions, we stained Cav1 KO and WT A431 cells with Di4 ANEPPDHQ that reports on membrane fluidity (as measured by generalized polarization (GP) of Di4 ANEPPDHQ) (Fig. 4J). For both control and Cav1 KO cells we observed a lower membrane fluidity (higher GP value) in the sorting endosome compared to the sorting tubules emerging from this compartment (Fig. 4J). But as predicted from our earlier results, the endosomal vesicles in the Cav1 KO cells displayed a significantly lower membrane fluidity compared to endosome vesicles in WT cells -consistent with a higher SL and cholesterol content. And correspondingly, we found a higher membrane fluidity (lower GP value) in endosomal sorting tubules of the Cav1 KO compared to endosomal tubules of WT cells – consistent with lower SL and cholesterol content (Fig. 4J).

These results altogether provide evidence that complex sphingolipids fail to enter recycling tubules emerging from the sorting endosome in Cav1 KO cells. Instead, they are retained in the endosome vesicle for maturation into the late-endosome/lysosome pathway.

### Endosome sorting of the sphingolipids into recycling tubules requires palmitoylation of Cav1

To understand how Cav1 may affect the endosomal sorting of complex SLs, we first defined the localization of Cav1 within the endosomal system using Cav1 KO cells (Suppl. Fig. 5A, B and Fig. 5A). Transient over-expression of Cav1-mCherry in A431 Cav1 KO cells showed an apparent close colocalization with internalized Alexa-labeled Tfn in both the sorting endosome and in recycling tubules emerging from these structures (Fig. 5A and Suppl. Fig. 5A). In MLEC Cav1 KO cells, transient over-expression of the endosome recycling tubule associated kinesin Kif13A showed co-localization with co-expressed Cav1-RFP (Suppl. Fig. 5B). These results are consistent with Cav1 localization to the endosome network in other experimental systems (Gagescu et al., 2000; He et al., 2015; Lapierre et al., 2012), including in cells lacking Cavin-1 (Verma et al., 2010). As such, we conclude Cav1 is likely positioned to act directly on the complex sphingolipids for sorting between the endosomal recycling and late-endosome/lysosomal pathways.

**Figure 5.**
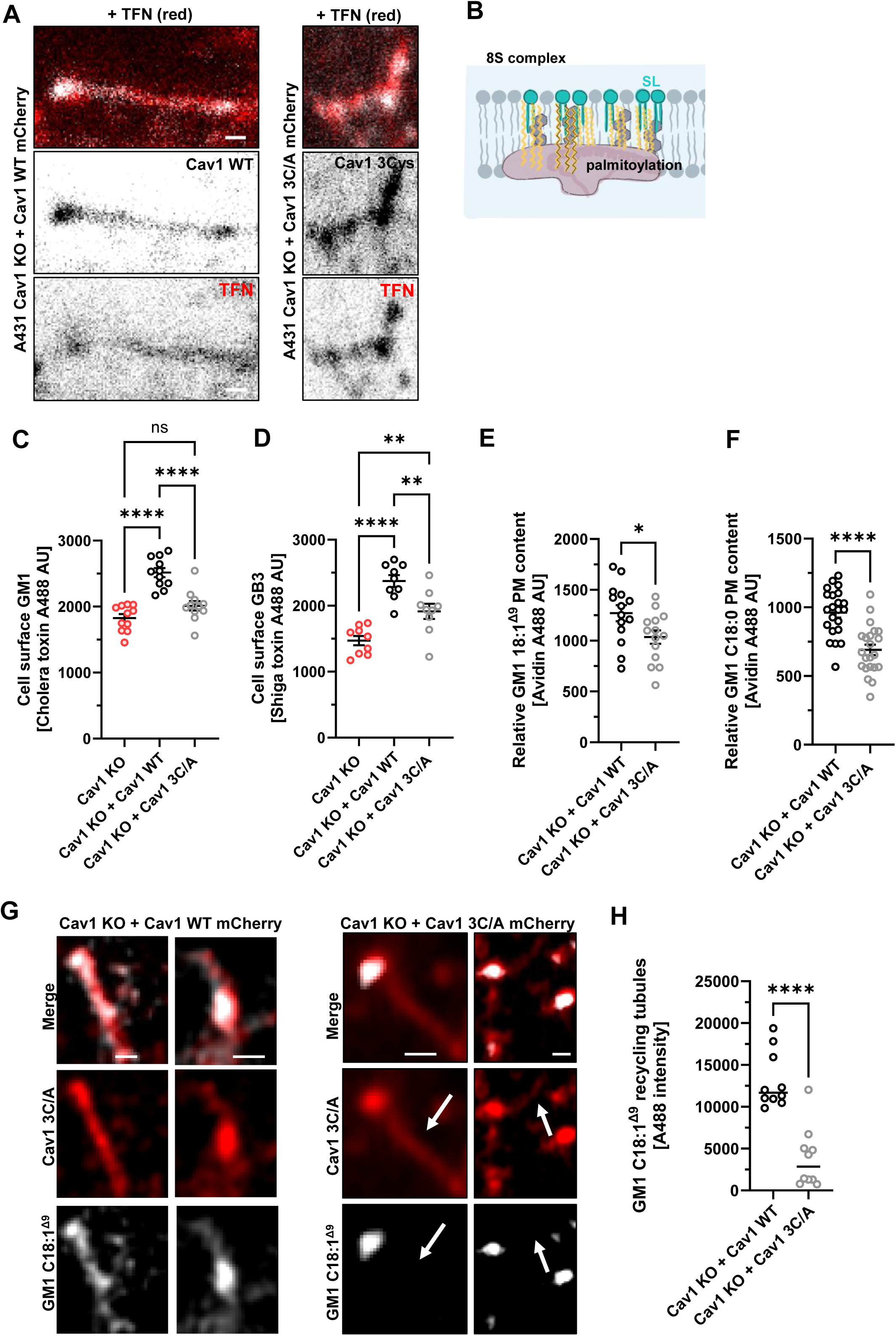
Palmitoylation sites in Cav1 are required for endocytic SL sorting. **(A)** Subcellular localization Cav1-mCherry and Cav1-3C/A mutant-mCherry to TFN positive recycling endosomal tubules in A431 Cav1 KO reconstituted with Cav1-mCherry or Cav1-3C/A mutant protein. Scale bar = 0.5 µm. **(B)** Cartoon displaying 8S complex of Cav1 (burgundy) with palmitoylations (yellow) reaching the outer membrane leaflet to interact with SL (turquoise). **(C)** Quantification of GM1 plasma membrane content by CTxB-A488 FACS in A431 Cav1 KO, Cav1 KO overexpressing Cav1 WT or Cav1-3C/A mutant mCherry protein. Mean and SEM displayed. N = >4 biological replicates, dots represent 20.000 cells. **(D)** Quantification of GB3 plasma membrane content by STxB-A488 FACS in A431 Cav1 KO, Cav1 KO overexpressing Cav1 WT or Cav1-3C/A mutant mCherry protein. Mean and SEM displayed. N = >4 biological replicates, dots represent 20.000 cells. **(E)** GM1 C18:1^Δ9^ plasma membrane recycling measured by streptavidin-A488 in A431 Cav1 KO overexpressing Cav1 WT or Cav1-3C/A mutant mCherry protein. Mean and SEM displayed. N = > 5 biological replicates, dots represent 20.000 cells. **(F)** GM1 C18:0 plasma membrane recycling measured by streptavidin-A488 in A431 Cav1 KO overexpressing Cav1 WT or Cav1-3C/A mutant mCherry protein. Mean and SEM displayed. N = > 5 biological replicates, dots represent 20.000 cells. **(G)** In A431 Cav1 KO cells overexpressing Cav1 WT mCherry, GM1 C18:1^Δ9^ co-localizes to Cav1 WT mCherry or TFN positive endosomal vesicles and tubules. In A431 Cav1 KO cells overexpressing Cav1 3C/A mutant mCherry, GM1 C18:1^Δ9^ co-localizes to the Cav1 3C/A mutant mCherry or TFN positive endosomal vesicles but not the tubules (white arrow). Exemplary images, scale bar = 0.5 µm. **(H)** Quantification of (G) for n = 3 biological replicates. Mean displayed. For all experiments: n.s. p > 0.05, *p < 0.05, **p < 0.01 ***p < 0.001 and ****p < 0.0001.

A recent cryoEM structure revealed that Cav1 forms an assembly of 11 protomers (the Cav1 8S complex) arranged in a symmetric spiral disc. The 8S complex is predicted to integrate within and to occupy the entire inner (cytosolic facing) lipid-leaflet of cell membranes (Fig. 5B) (Porta et al., 2022). Each Cav1 monomer contains 3 cysteine residues that are substrates for S-palmitoylation (Dietzen et al., 1995; Parat and Fox, 2001). All are positioned within the newly defined spoke region on the flat surface of the Cav1 8S complex facing the opposing outer-membrane lipid-leaflet (Doktorova et al., 2025; Han et al., 2025; Porta et al., 2022). The 11 protomer complex can thus potentially bear as many as 33 palmitoylations positioned to interact with the outer-membrane leaflet lipids, including the complex sphingolipids (Doktorova et al., 2025; Liebl and Voth, 2024). As such, we hypothesized that the Cav1-palmitoylations of the 8S complex may act to chaperone entry of the complex sphingolipids into the narrow highly curved sorting tubules emerging from the sorting endosome. To test this, we generated a Cav1 mutant lacking all 3 palmitoylation sites by replacing the cysteine residues with alanine (Cav1-3C/A). When overexpressed in A431 Cav1 KO cells, we found the Cav1-3C/A-mCherry mutant protein still localized to Tfn-labeled sorting endosomes and recycling tubules, the PM, and lysosomes as observed for the WT Cav1 protein (with both WT and mutant expressed at similar levels) (Fig. 5A and Suppl. Fig. 5A and C).

We then asked if the Cav1-3C/A mutant could rescue the SL trafficking defects observed in Cav1 KO cells. While overexpression of WT Cav1-mCherry restored cell surface levels of endogenous GM1 and GB3 – as measured by cholera and Shiga toxin PM binding respectively (Fig. 5C and D and Suppl. Fig. 1F and G), this was not the case for cells expressing the Cav1-3C/A mutant protein (Fig. 5C and D). When plasma membrane recycling for the biotin-labeled GM1 C18:0 and GM1 C18:1 species was measured, we found significantly less GM1 in the PM over time after pulse-chase for both GM1 species in Cav1 KO cells expressing the Cav1-3C/A mutant -as compared to cells expressing WT Cav1 (Fig. 5E and F). Furthermore, overexpression of Cav1 WT protein, but not mutant Cav1-3C/A, in Cav1 KO cells rescued the sorting of GM1 C18:1 into recycling endosomal tubules (Fig. 5G and H); and this reduced the high membrane order (higher GP value) in the sorting vesicles (Suppl. Fig. 5D). These results show the Cys residues are required for endosomal sorting of GM1 when assisted by Cav1, and implicate palmitoylation as a causal factor.

### The cysteine palmitoylation sites within the spoke region of Cav are conserved

We analyzed the degree of conservation of the three cys residues in human Cav1 on 74 evolutionarily diverse caveolins recently studied (Han et al., 2025). Caveolins are found throughout Metazoa and are also present in the choanoflagellate *S. rosetta* (Han et al., 2025). Here, we focused on the number and positioning of cysteines mainly within the spoke region (residues 109-169) of human Cav1 (Han et al., 2025). Notably, Cys positions C133, C143, and C156 seen in human Cav1 did not appear as the most prominent potential palmitoylation sites in caveolins across metazoan evolution (Fig. 6A). They are concerved throughout the chordates with the exception that the three *Xenopus* Cav1 homologs lack Cys at the equivalent residues 143 and 156 (Fig. 6A and Suppl. Fig.6A). The three Cys residues in human Cav1 are however highly conserved across all vertebrates in Caveolin-3, a paralogue that like Cav1 assembles with Cavin-1 to form caveolae (Suppl. Fig. 6B) (Bastiani et al., 2009). Cav3 was also predicted to assemble into a disc-shaped structure, similar to the Cav1 8S complex, a prediction supported by observations with negative-stain electron microscopy (Han et al., 2025; Whiteley et al., 2012) Recent studies show that Cav3 functions on the highly curved T-tubules of muscle myocytes (analogous in geometry to endosome sorting tubules) to shape their biology (Lemerle et al., 2023).

**Figure 6.**
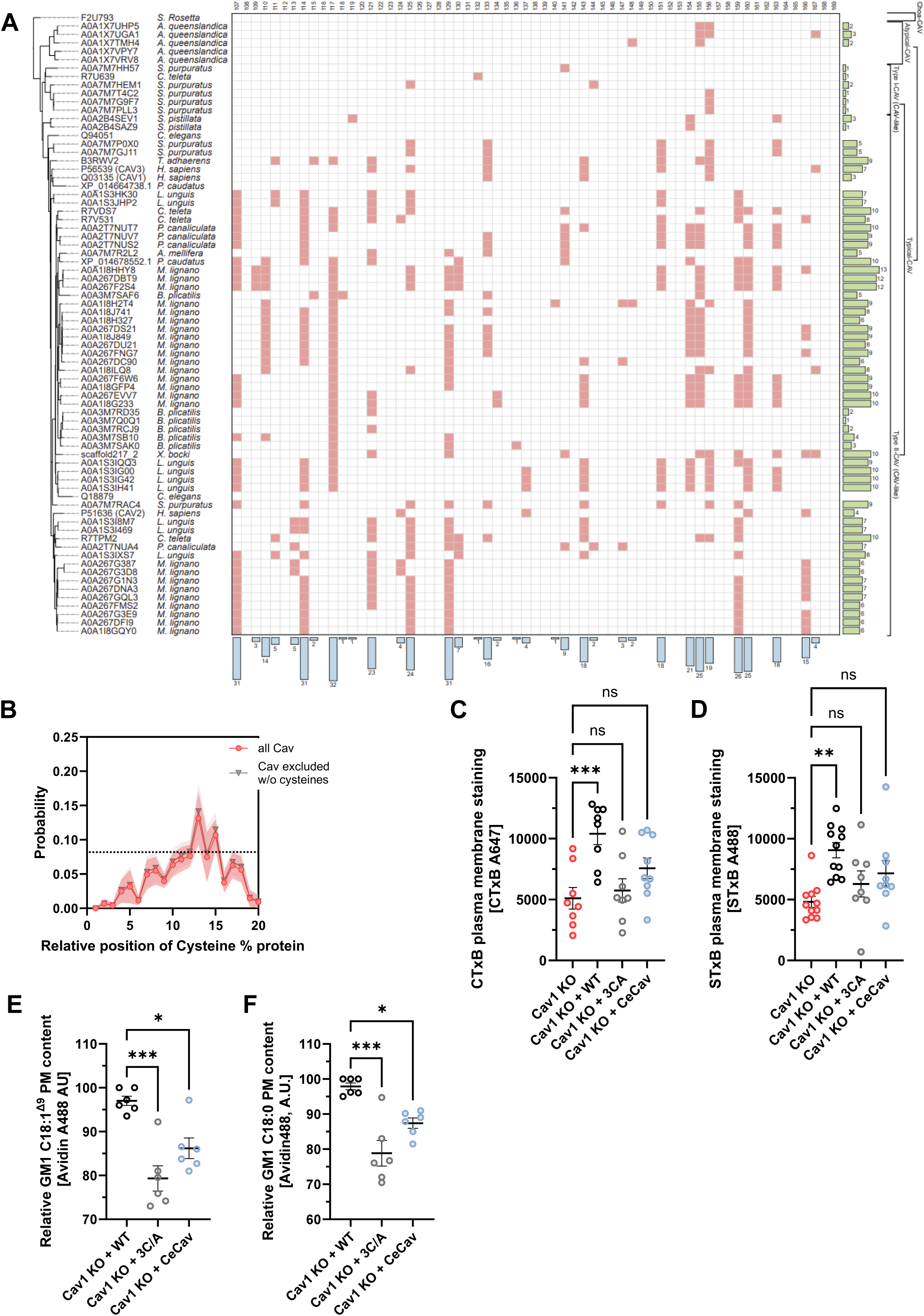
Evolutionary analysis of three palmitoylation sites within the Cav1 spoke region. **(A)** Evolutionary analysis of potential palmitoylation sites on cysteine residues across 74 caveolin paralogues within the predicted spoke region (amino acids 107-169 in human Cav1). Total cysteine numbers in green bars, cysteine sites are highlighted in red. All but six sequences contain cysteines within the spoke region. **(B)** Probability analysis for cysteine to occur across the amino acid sequence of caveolin paralogues including (red) and excluding (grey) paralogues that lack cysteine within the spoke region. **(C)** Cell surface amount of GM1, measured by FACS using CTxB-A647 in A431 Cav1 KO or Cav1 KO overexpressing Cav1 WT, Cav1-3C/A-mCherry or CeCav. Mean and SEM displayed. N=> 3 biological experiments, dots represent 20.000 cells. **(D)** Cell surface amount of GB3, measured by FACS using STxB-A488 in A431 Cav1 KO or Cav1 KO overexpressing Cav1 WT, Cav1-3C/A- mCherry or CeCav-myc. Mean and SEM displayed. N=> 3 biological experiments, dots represent 20.000 cells. **(E)** Plasma membrane recycling assay after 1.5h for GM1 C18:1^Δ9^ by FACS in A431 Cav1 KO or Cav1 KO overexpressing Cav1 WT, Cav1-3C/A, or CeCav. N=> 3 biological experiments, dots represent 20.000 cells, mean and SEM displayed. **(F)** Plasma membrane recycling assay after 1.5h for GM1 C18:0 by FACS in A431 Cav1 KO or Cav1 KO overexpressing Cav1 WT, Cav1-3C/A-mCherry or CeCav-myc. N=> 3 biological experiments, mean and SEM displayed, dots represent 20.000 cells. For all experiments: n.s. p > 0.05, *p < 0.05, **p < 0.01 ***p < 0.001 and ****p < 0.0001.

While the exact number and positions of the cysteine residues in the spoke region of caveolins were not strictly conserved across the evolutionary tree we studied, when compared to the average frequency of cysteine residues in polypeptide sequences in proteins (*H. sapiens* 2.3%, *D. melanogaster* 1.9 %, archaea 0.5%) (Castillo-Villanueva et al., 2023; Wiedemann et al., 2020) we noted a conserved enrichment of cysteines within the spoke region (Fig. 6B). Only six caveolin proteins lacked at least 1 cysteine in this region - *S. rosetta* (F2U793), two of the five paralogues in *A. queenslandica* (A0A1X7VPY7 and A0A1X7VPY8), *P. caudatus* (XP_014664738.1), and the two *C. elegans* caveolins (Q94051 and Q18879). We also noted a depletion of cysteine residues in the first 60 to 80 amino acids of caveolin in all species analyzed (Fig. 6B). Thus, in almost all cases, the caveolins appear to have a unique need for concentration of cys residues in the spoke region to accommodate protein function.

The naturally occurring caveolins lacking all cysteines in the spoke region provided an experiment of nature to test the palmitoylation hypothesis. As such, we studied a non-human caveolin for its ability to sort the sphingolipids into endosome recycling tubules in human A431 Cav1 KO cells - the invertebrate caveolin from *C. elegans* (CeCav1/CeCav-a, Q94051). This protein lacks cysteine residues within the spoke region, but is predicted to assemble as an 11-mer 8S complex by AlphaFold (Suppl. Fig. 6B) (Han et al., 2025). It also was shown to be expressed diffusely on the plasma membrane when expressed in Cav1^-/-^ MEFs or Hela cells (Kirkham et al., 2008) (Kenworthy unpublished).

*C. elegans* CeCav1 was transiently expressed in Cav1 KO A431 cells and plasma membrane levels of endogenous GM1 and Gb3 were measured by FACS using fluorescent Cholera and Shiga toxin respectively (Fig. 6C and 6D). We also measure plasma membrane levels of exogenously added GM1 C18:1 and GM1 C18:0 after pulse-chase (Fig. 6F and G) . We found that, unlike WT Cav1, CeCav1 failed to rescue the defects in PM content or apparent endosome recycling for these complex sphingolipids. Rather, this *C. elegans* paralogue phenocoped our results with mutant human Cav1 lacking the three cys residues in the spoke region. These results strengthen the evidence that palmitoylation of the spoke region underlies the ability of human Cav1 to enable endosome sorting of the complex sphingolipids.

## DISCUSSION

The results of these studies show that Cav1 acts independently of caveolae to enable the sorting of the complex sphingolipids into endosomal recycling tubules, thus explaining in part how eukaryotic cells traffic and maintain the content of complex sphingolipids in the plasma membrane. This is a newly appreciated function for caveolin-1. For biophysical and mechanical reasons, it was found unfavorable for the sphingolipids and cholesterol to become enriched into narrow and highly curved tubules emerging from liposomes (Safouane et al., 2010; Sorre et al., 2009; Tian and Baumgart, 2009). Thus, purely physical mechanisms of sphingolipid demixing and nanodomain formation with differential affinity for membrane curvature are predicted to be insufficient to account for the presence of sphingolipids in endosome sorting tubules of live cells. In vitro studies using giant liposomes indicate that the enrichment of complex sphingolipids into curved structures required direct or indirect interaction with membrane or membrane-associated proteins that recognize membrane curvature and assiste the recruitment of these lipids into tubules formed from the originating liposome (Safouane et al., 2010; Sorre et al., 2009). This prompted our interest in Cav1, a membrane protein distributed across a variety of cell surface and intracellular membranes of the secretory and endocytic pathways. Cav1 interacts strongly with sphingolipids and cholesterol and has been previously implicated in the sorting of these lipids - but with mechanism largely unknown (Cheng et al., 2006; Lemerle et al., 2023; Parker et al., 2009; Pol et al., 2020; Pol et al., 2001; Shvets et al., 2015). Here we have tested this hypothesis by measuring the trafficking of two complex sphingolipids, sphingomyelin and the glycosphingolipid GM1, and their dependence on Cav1 for proper trafficking and subcellular distribution.

We found that Cav1 affected the sorting of the complex sphingolipids into endosomal recycling tubules. Cav1 acted on the complex sphingolipids independently of Cavin-1 and thus independently of caveolae. Such caveolae-independent functions of Cav1 may also occur in lymphocytes and neurons which express Cav1 but lack expression of the cavins, and in the pre-vertebrates where Cav1 is also expressed but not the cavins (Han et al., 2025; Hansen and Nichols, 2010). Our results are also consistent with observations of caveolae-independent functions of Cav1 in other cell types in vertebrates (Lolo et al., 2023; Minguet et al., 2017; Shikanai et al., 2018).

The effects of Cav1 on the complex sphingolipids likely also applies to their sorting in the secretory pathway. Evidence for this was more definitively found in our studies using the model GPI-anchored protein GPI-GFP. Similar results were recently observed for trafficking nascent GPI- anchored proteins through the Golgi complex in cells lacking Cav1 (Carpentier et al., 2025). In those studies, using an equivalent GPI-RUSH system as in our study, Carpentier et al. showed that in cells lacking the lipid droplet protein seipin 1, the redistribution of Cav1 from the secretory pathway to lipid droplets caused depletion of GPI anchored proteins in the plasma membrane, associated with an increase in membrane fluidity in the TGN/secretory compartments. Both observations are consistent with our results and with earlier studies in cells lacking Cav1 (Sotgia et al., 2002).

The ability of Cav1 to chaperone the complex sphingolipids required its palmitoylation. Palmitoylation likely enhanced the strength of association between Cav1 in the inner membrane leaflet with the sphingolipids located in the outer leaflet. The three palmitoylation sites, the Cys residues C133, C143, and C156, locate to the spoke region of Cav1 which directly interfaces with the outer membrane lipid leaflet where the complex sphingolipids reside. Recent atomistic and coarse-grained simulations predict the Cav1 8S complex to associate with positively curved membranes, a characteristic of endosome sorting tubules. These studies also showed that the 8S complex can affect the composition of the outer membrane leaflet by attracting between 40-70 cholesterol molecules into the region adjacent to the Cav1 8S complex (Doktorova et al., 2025). Models of the 8S complex fully palmitoylated showed a greater effect on cholesterol recruitment and cholesterol ordering (Doktorova et al., 2025; Liebl and Voth, 2024).

Given the strength of association between the complex sphingolipids and cholesterol (London, 2005; Ramstedt and Slotte, 1999; Ramstedt and Slotte, 2006), such an enrichment of cholesterol could provide a driving force for sequestration of the complex sphingolipids in that region of the opposing membrane leaflet adjacent to the 8S complex. This may explain how Cav1 associates with these sphingolipids (and cholesterol) and enable their entry into the highly curved endosome sorting tubules that shape PM structure and function. Both GM1 species studied were affected by palmitoylated Cav1 – GM1 C18:0 that contains the C14* motif enabling tight association with cholesterol and assembly in nanodomains, and also GM1 C18:1 that does not. The affinity for GM1 C18:1 to associate with the Cav1-induced cholesterol enriched region is likely explained by its residual but still significant affinity for cholesterol due to hydrogen-bonding between the cholesterol’s hydroxyl headgroup and the amide on the sphingosine chain – enhanced by the extensive hydrophobic interactions between cholesterol and the rest of the ceramide molecule (Ramstedt and Slotte, 1999; Slotte, 2016).

The three cysteines of Cav1 (C133, C143, and C156) were found to be evolutionarily conserved across the vertebrate clades of Cav1 and Cav3. These cysteine residues in Cav1, however, were not conserved throughout all the metazoan clades examined – though we found an enrichment of cysteine residues within the spoke region of the caveolins dating back to the sponge ponifera clade *A. queenslandica*. In the case of Cav3, the three cysteine residues were more highly conserved, appearing in all clades well before vertebrate evolution. Furthermore, human Cav3 contains an additional three cysteines within this region, resulting in up to six cysteines that can be palmitoylated (Ashford et al., 2024). We find this notable as Cav3 operates at the juncture of the highly curved T-tubules of myocytes – enabling the selective trafficking of lipids and proteins into these structures – consistent with how Cav1 appears to function in the structurally analogous endosome sorting tubules of the vertebrates (Lemerle et al., 2023). Such a chaperone-like activity may represent an emergent function of the Cav1 8S complex when present on highly curved membranes of endosome sorting tubules or tubules of the clathrin-independent CLIC pathway (or in spherical caveolae buds when associated with cavin1 – but then facilitating the exchange of the complex sphingolipids when the Cav1 8S - sphingolipid complexes move onto flat membranes of the endosomal compartment and PM. This work defines a newly discovered form of lipid sorting dependent on Cav1 and affecting membrane structure and function.

## Acknowledgements

We thank the Harvard Center for Biological Imaging, Boston Children’s Hospital Cell Function and Imaging Core, and HDDC Imaging Core for infrastructure and support. We acknowledge the Cell and Tissue Imaging Core facility (PICT IBiSA), Institut Curie, member of the French National Research Infrastructure France-BioImaging (ANR10-INBS-04). We further thank Dr. Krishnan Raghunathan and Dr. Ajit Tiwari for helpful discussions during the preparation of this manuscript, Dr. Darrin T. Schultz for advice regarding the phylogenetic analysis, and Yelena Peskova for expert technical assistance.

This work was supported by NIH grants DK048106, DK122953, DK118640, and the P30 Harvard Digestive Disease Center DK034854 to W.I.L., a Charles A. King Trust Postdoctoral Research Fellowship and a Boston Children’s Hospital Faculty Career Development grant to S.S.S., and R01HL168258 and R01GM151635 to AKK. JP, PB and CL were funded by the grant ANR-20-CE13-0002 from the Agence Nationale pour la Recherche. PB team is supported by the Fondation pour la Recherche Médicale (FRM) (FRM EQU202003010307). PB is member of the CNRS consortium AQV. JP, PB and CL are members of the Labex Cell(n)Scale (ANR-11-LABX0038) and Paris Sciences et Lettres (ANR-10-IDEX-0001-02).

The content is solely the responsibility of the authors and does not necessarily represent the official views of the National Institutes of Health.

## Author contributions

S.S.S. performed the cell biology, lipid chemistry, lipidomics and imaging experiments on A431 cells. J.P. performed the cell biology, lipid chemistry, lipidomics and imaging experiments on MLEC cells. C.V. assisted with MLEC experiments. L.H. assisted with Cav1 3C/A experiments. N.M. assisted with lipidomics analysis. R.T. performed lipidomics. B.H. performed evolutionary analysis. S.S.S., J.P., H.B., W.I.L., C.L., P.B. and A.K.K. wrote the manuscript. W.I.L., C.L., P.B. and A.K.K. supervised and directed the research. All authors discussed the manuscript and contributed to the preparation of the manuscript.

## Material and Methods

### Reagents

Sphingomyelin (SM), Cholesterol (ChL) and DOPC were purchased from Avanti Research, DHPE-Texas Red and DAPI from Thermo Fisher. D-biotin, Dimethylformamide (DMF), dimethylsulfoxide (DMSO), defatted bovine serum albumin (dfBSA), fetal bovine serum (FBS), and filipin III (filipin) were purchased from Sigma Aldrich (St. Louis, MO). Dulbecco’s modified Eagle’s medium (DMEM) with or without phenol red, OptiMEM®, Lipofectamine® 2000, Laurdan (6-Dodecanoyl-2-Dimethylaminophthalene), *FAST*Dil™ Dil^Δ9,12^-C_18_(3), Di-4-ANEPPDHQ, AlexaFluor™ (−405, −488, −568, or −647 nm) and pHrodo™ Fluor -NHS ester and –azide dyes, AlexaFluor™ and pHrodo™ Fluor conjugates, Dextran Pacific Blue™ (4000 MW) and LysoTracker™ were purchased from Thermo Fisher Scientific (Waltham, MA). Cholera toxin B subunit and Shiga toxin B subunit were purchased from Sigma Aldrich (St. Louis, MO).

### Primers

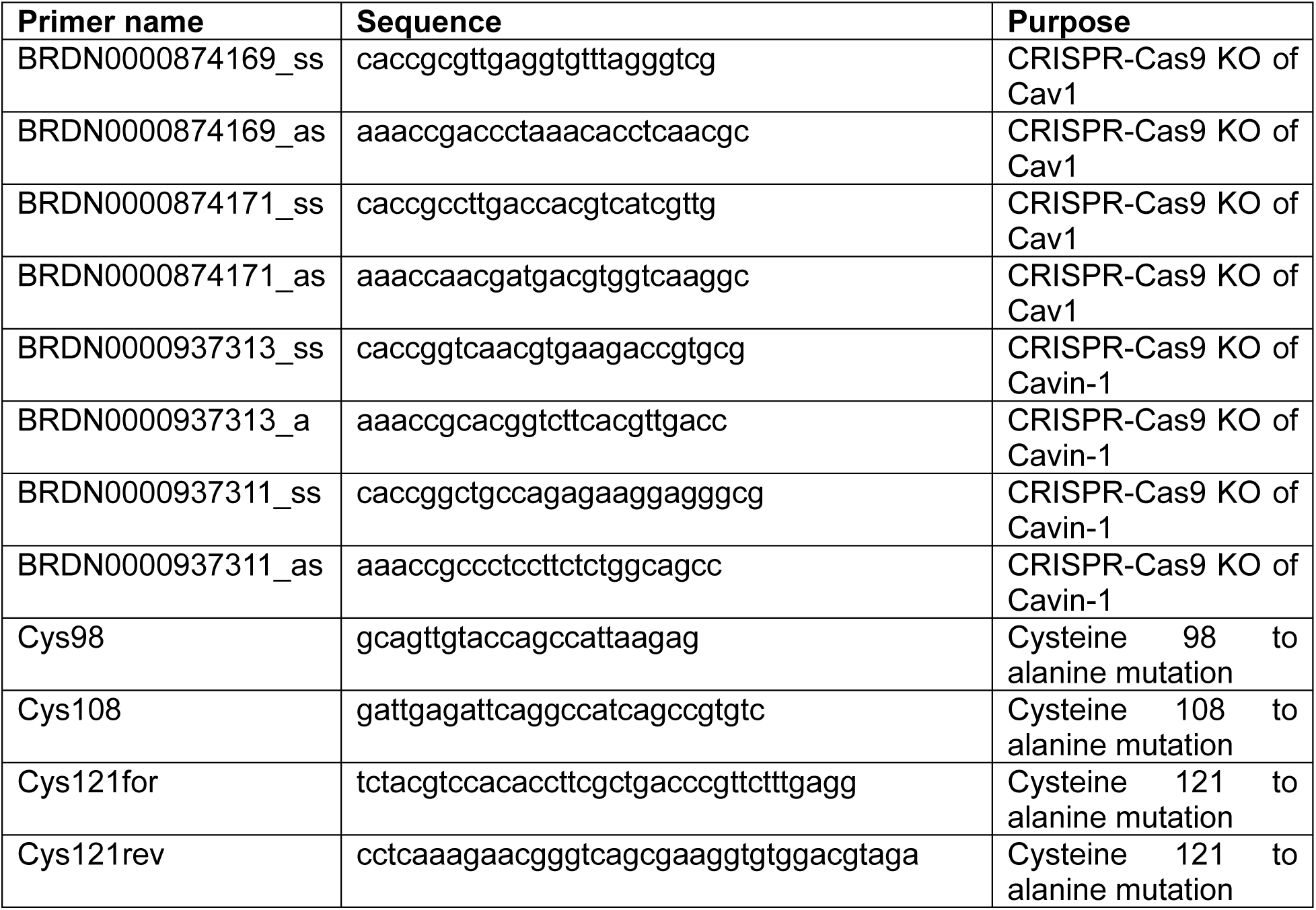
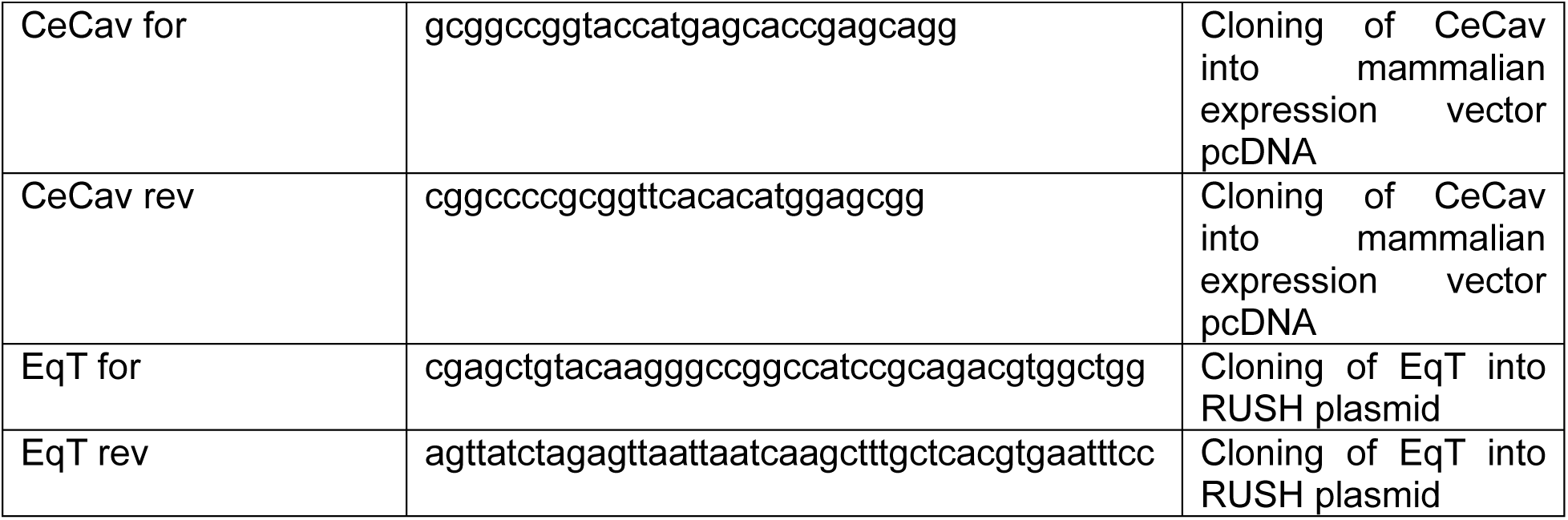

### Plasmids

pET28a_EqT plasmid kind gift from Christopher G Burd

Str-KDEL_SBP-NPY and GPI-GFP plasmids kind gift from Franck Perez

Str-KDEL_SBP-EqT this study

Str-KDEL_SBP-EqTsol this study

pDONR201_Kif13A plasmid kind gift from Cedric Delevoye

Addgene plasmid lentiCas9-Blast (Addgene #52962)

Addgene plasmid lentiGuide-Puro (Addgene #52963)

pVSVg and psPAX2 (Addgene plasmids #8454, #12260)

pcDNA_Caveolin-1_mRFP kind gift from Richard Pagano

pcDNA_Caveolin-1_mCherry (Hanson et al., 2013)

pcDNA_Caveolin-1_3C/A_mCherry this study.

pcDNA_CeCaveolin_myc this study.

### Antibodies

Caveolin-1 (Cell Signaling, #3238S), Cavin-1 (Proteintech, #18892-1-AP), SPTLC-2 (Abcam, AB236900), Ceramide synthase 5 (Sigma, SAB4301211), Ceramide synthase 2 (Thermo, CF809918), UGCG (Thermo, BS-8593R), B4GALT4 (ATLAS Antibody, HPA063546), Lamp1 (Abcam ab24170).

### Cell lines and culture conditions

A431 and HEK293T cells were purchased from American Type Culture Collection, VA and maintained in DMEM (Gibco) with 10% FBS (Gibco) and 1% Pen/Strep (HyClone). MLEC cells were maintained in EBM2 media (Lonza) supplemented with EGMTM-2 Endothelial Cell Growth Medium-2 BulletKit (Lonza), 10% FBS (Gibco) and 1% Pen/Strep (Gibco).

Cells were routinely split at 80% confluency and plated for experiments 48 hours prior experiments. Cell density at the time of microscopy was 70% confluent. Cells were transfected at 50 % confluency with respective plasmids using Lipofectamine 3000 reagent and 750ng plasmid DNA/12well. Transfected cells were used 16 h post transfection. MLEC were routinely split at 80% confluency and plated for experiments 24 hours prior experiments. Cell density at the time of microscopy was 70% confluent. MLEC cells were transfected using electroporation with Ingenio Electroporation Solution (MirusBio). Transfected cells were used 16h post transfection, unless stated differently.

### Generation of EqT-RUSH construct

Eqt-SM and Eqt-Sol were cloned into plasmid Str-KDEL_SBP-EGFP-Ecadherin (Addgene #65286). Eqt was cloned in the place of Ecadherin with FseI and Pac1 restriction sites.

### Generation of Caveolin-1 with no palmitoylation sites

Site directed mutagenesis for Cys 98, Cys 108 and Cys 121 was performed using Agilent QuickChange and QuickChange Lightening site-directed mutagenesis kit on pcDNA_Caveolin-1-mCherry template and primer pair C98Afor and C108Afor. The resulting plasmid pcDNA_Caveolin-1-mCherry with C98A/C108A was used as a template with the primer pair C121Afor and C121Arev and Agilent QuickChange to generate pcDNA_Caveolin-1-3C/A. All plasmids were verified by sequencing.

### Generation of Caveolin-1 with no palmitoylation sites

The gene encoding the entire caveolin polypeptide of *C. elegans* caveolin (Q94051) was synthesized by Genscript, NJ. It was amplified by the PCR using PfuUltra II Fusion HS DNA Polymerase (Agilent, #600670) to introduce KpnI and SaII restriction sites with primers CeCav for and CeCav rev. The digested PCR product was subsequently cloned into the modified pcDNA backbone with the C-terminal myc-tag generated by KpnI and SacII (New England Biolabs, MA) digest from the H. sapiens Caveolin1-myc plasmid described in (Hanson et al., 2013). SacII site was destroyed by the mutagenesis using Quick Change Lightning kit (Agilent, #210518) Primers used were: 5’-CGCTCCATGTGTGAAGCTTGGGCCCGAAC-3’(sense) and 5’-GTTCGGGCCCAAGCTTCACACATGGAGCG-3’(antisense). The resulting plasmid was verified by Sanger sequencing (Genewiz/Azenta Life Sciences, NJ).

### Generation of CRISPR-Cas9 KO cell lines

Generation of an A431 cell line stably expressing Cas9 and the CRISPR-Cas9 mediated knock-outs for Caveolin-1 and Cavin-1 were generated using lentiCas9-Blast for stable integration of Cas9 endonuclease. Guide sequences against Caveolin-1 and Cavin-1 were cloned into lentiGuide-Puro (see Table 1). Plasmid integrities were verified by sequencing (IDT technologies, Coralville, IA). Lentivirus was produced by co-transfecting HEK293T cells with packaging plasmids pVSVg and psPAX2 in the presence of polyethylenimine (PEI). 48h post-transfection supernatants containing virus were collected and purified using 0.2µm. A431 Cas9 cell line was incubated with respective virus for 48h, subsequently, cells were selected for 7 days in the presence of Blasticidin and/or Puromycin. CRISPR-Cas9 KO cell lines were verified by lysing the cells and immunoblotting for Caveolin-1 or Cavin-1.

### Protein Expression and Purification of EqT

The EqT (SM or Sol) plasmid was transformed into BL21 (DE3) - RIPL cells. The cells were grown overnight in 50 mL of 2YT media containing kanamycin and chloramphenicol at 37°C under agitation. The following day, cells were diluted into 2 L of 2YT media with antibiotics and grown in a shaker incubator at 37°C to an optical density at 600 nm (OD600) of ∼0.6. Expression of the protein was induced by adding 0.4 mM IPTG, and cells were incubated for 4 hours at 37°C with shaking.

Subsequently, cells were harvested by centrifugation at 10,000 × g for 10 minutes. The supernatant was discarded, and the cell pellet was resuspended and washed with PBS. The cell pellet was then resuspended in 40 mL of wash buffer (20 mM Na₂HPO₄/NaH₂PO₄, pH 7.4, 500 mM NaCl, 25 mM imidazole), supplemented with Complete Mini EDTA-free protease inhibitor and 1 mM PMSF, and lysed by sonication for 10 minutes with 3-second on and 3-second off pulses.

The lysate was centrifuged at 20,000 × g for 30 minutes at 4°C, and the supernatant was collected. The cleared extract was applied to Ni²⁺-NTA beads (Thermo Fisher Scientific) and incubated with agitation for 2 hours at 4°C. The beads were washed three times with 50 mL of wash buffer using the batch method. The beads were then packed into a column and washed an additional 10 column volumes (CV) with the same buffer.

Protein was eluted with 10 mL of elution buffer (20 mM Na₂HPO₄/NaH₂PO₄, pH 7.4, 500 mM NaCl, 300 mM imidazole). The eluted protein was concentrated to ∼500 µL using a 10 kDa concentrator and further purified by FPLC on a Superdex Increase 200 10/300 GL column with 20 mM Na₂HPO₄/NaH₂PO₄, pH 7.4, 500 mM NaCl buffer. Fractions containing EqT were combined, and the protein concentration was determined by Bradford assay.

For labeling, EqT was conjugated with Alexa Fluor™ 647 (or 488) NHS Ester (Invitrogen) at a 1:1 molar ratio with overnight agitation at 4°C. Free dye was removed by dialysis against 20 mM Na₂HPO₄/NaH₂PO₄, pH 7.4, 500 mM NaCl buffer, followed by concentration to ∼500 µL using a 10 kDa concentrator. To preserve the labeled protein, 0.02% sodium azide was added. Labeled proteins were stored at 4°C for up to 3 months.

OlyA-His6-WT overexpression was induced at 18°C with 1 mM IPTG for 16 h, and the recombinant protein was purified by nickel chromatography followed by gel filtration (Platform of protein production, Institut Curie) using the same procedures as described in (Endapally et al., 2019). After purification, OlyA-His6 was labeled with Alexa Fluor 647TM C2-maleimide dye as described previously (Endapally et al., 2019).

### Synthesis of peptide and fluorophore-labeled GM1 species

GM1 C18:1 was supplied by Prof. Sandro Sonnino (U. Milan, Italy). Peptide containing modified functional residues were custom synthesized by Biosynth (Gardner, MA) as all-D peptide with the following amino acid sequence: propargylglycine-k(ε-biotinoyl)(ds)g(dy)g(dr)g(ds)g-(kaoa)- amine. Synthesis of peptide-lipid and fluorophore-labeled GM1 conjugates was performed using previously published work from our lab (Arumugam et al., 2021; Chinnapen et al., 2012; Garcia-Castillo et al., 2018; Schmieder et al., 2022; te Welscher et al., 2014). In brief, GM1 species were oxidized with sodium periodate and purified over Bond Elute SepPak C18 cartridges using methanol as eluent (Agilent, MA). Oxidized GM1 species were reacted in a 1:1.5 ratio with functional aminooxy-containing peptides and products were purified by HPLC and verified by LC-MS. GM1-peptide fusions were labeled with AlexaFluor-azide via copper mediated click chemistry. Compounds were purified by HPLC and verified by mass spectrometry. Working GM1 stock solutions were prepared in 33% DMF/ddH20.

### RUSH assay live imaging

MLEC cells were transfected via electroporation with the indicated RUSH and Mannosidase II plasmids and seeded onto an 8-well imaging glass slide chamber (Ibidi) 24 hours prior to imaging. The following day, the slides were placed on the stage of a microscope equipped with a temperature-controlled, CO₂-buffered chamber. Media was gently aspirated and replaced with fresh media containing 40 μM D-biotin (Sigma-Aldrich). Time-lapse imaging was performed at 37°C using an Eclipse 80i microscope (Nikon) with a spinning disk confocal head (PerkinElmer) and a CoolSnapHQ2 camera (Roper Scientific). For Golgi exit dynamics, images were captured with a 40x objective using MetaMorph software (Molecular Devices) at 2.5-minute intervals for approximately 90 minutes. Full Z-stacks were acquired with a 0.5 μm step size. Integrated intensity was quantified at each time point in a region of interest (ROI) encompassing the Golgi complex, which was defined by the Mannosidase II fluorescent signal. These values were normalized to the maximum value. Image analysis was conducted using ImageJ software, and plots were generated in GraphPad Prism. For Golgi-exiting vesicle dynamics and size, images were acquired with a 63x objective at 2-second intervals for approximately 1 minute, and only three central sections with a 0.5 μm step size were captured.

For cell surface delivery of EqT, MLEC cells were transfected and seeded onto 8-well imaging glass slide chambers pre-coated with anti-GFP antibody. Briefly, glass slides were incubated with 0.1 M carbonate buffer (pH 9.5) for 1 hour at 37°C, followed by a wash and incubation with 0.01% poly-L-lysine for 1 hour at 37°C. The slides were then washed with PBS, air-dried, and immersed in 50 µg/mL rabbit polyclonal anti-GFP antibody (Institut Curie Platform) in carbonate buffer for overnight incubation at 4°C. The following day, before cell seeding, slides were washed in PBS, and transfected cells were added to the chambers. Cells were allowed to grow for 24 hours. The next day, the chambers were placed on the microscope as described above. Upon media change to include biotin, images of the central slice were taken at 2-minute intervals to determine the time when EqT reached the Golgi, marking time 0. Subsequently, WGA-Ax647 at 5µg/ml (Thermo Fisher) was added to the media to label the cell surface, and images of 2-3 bottom sections (0.5 µm step size) were acquired every 5 minutes. EqT fluorescent signal intensity was quantified over time using ImageJ software on Z-sum projection images.

Data was exported to Prism and area under the curve was calculated for time points 0 to 60 min. Data points represent 2-3 biological experiments with ≥ 3 cells analyzed.

### Analysis of postGolgi EqT vesicles

Images acquired for RUSH were thresholded based on EqT intensity using Otsu threshold in ImageJ. The resulting binary images were analyzed for vesicle size and number. Vesicle size was limited to 10 to 100 pxls, size and number were extracted. To analyze speed and traveling distance for EqT positive vesicles, TrackMate plugin for ImageJ was used, the vesicle size was restricted again to 10 – 100 pxls and velocity as pxl/sec and average distance as px were extracted.

### Lysosomal colocalization of EqT vesicles

EqT RUSH cells were co-labeled with lysotracker to demarcate the acidic lysosomal compartments. Colocalization analyses were conducted using a custom Matlab program. Images were binarized by adaptive thresholding with a sensitivity of 0.27. The binarized image were morphologically opened twice with disk structuring element first with a radius of 2 and decomposition of 0 and second with a radius of 2 and a decomposition of 4. In each image and each time point, the pixels that were non-zero in either channel were used to calculate the Pearson’s correlation between the two channels.

### Immunofluorescence

For all immunostainings MLEC cells were seeded onto glass coverslips 24 hours prior to staining. To stain cell surface SM, the following day, cells were incubated with 0.5 µM EqT-Alexa647 in PBS for 5 minutes at 37°C. After incubation, cells were washed with ice-cold PBS, fixed for 10 minutes in 4% paraformaldehyde (PFA), and the free aldehydes were quenched with 50 mM NH₄Cl for 10 minutes. Coverslips were mounted in Fluoromount-G (Fisher Scientific) supplemented with DAPI to stain DNA.

For intracellular staining, after fixation and aldehyde quenching, cells were permeabilized with 0.05% saponin in PBS containing 2 mg/mL bovine serum albumin (BSA) for 20 minutes. Cells were then incubated with 0.5 µM EqT-Alexa647 in saponin/BSA for 45 minutes. Coverslips were mounted in Fluoromount-G (Fisher Scientific) with DAPI for DNA staining.

For colocalization with lysosomes, following permeabilization with saponin, cells were incubated with anti-Lamp1 antibody (Abcam) at a 1:100 dilution in saponin/BSA, followed by incubation with 0.5 µM EqT-Alexa647 and donkey anti-rabbit Alexa488 (Invitrogen) at a 1:200 dilution in saponin/BSA.

For Cav1 rescue experiments, MLEC Cav1 KO cells were transfected via electroporation with the Cav1-RFP or RFP only plasmids. 72h after electroporation cells were fixed and stained as above.

All images were acquired using a Nikon A1RHD25 confocal microscope with a 40x CFI Plan Apo Lambda S Sil objective. Z-stacks of the entire cell were captured, with 0.25μm step, and image analysis was performed using ImageJ. Quantification was conducted on Z-sum projections. Manders colocalization coefficient of EqT signal in Lamp1 was determined on a single plane image using JACOP plugin (Bolte and Cordelières, 2006). Statistical analysis and graph preparation were carried out using GraphPad Prism.

### GUVs preparation and EqT staining

Giant unilamellar vesicles (GUVs) were prepared via electroformation on Indium Tin Oxide (ITO) plates. ITO plates were cleaned with ethanol and water, then dried under a nitrogen stream. A thin layer of chloroform was spread onto the ITO plate before lipid deposition to facilitate even spreading of lipid mixtures. Lipid mixtures were prepared from chloroform stock solutions and a small amount of the mixture was applied to the ITO plate using a Hamilton syringe. The lipid film was dried for 1 hour under vacuum. The ITO chambers were sealed, and 210 mM Sucrose solution was injected between the slides. GUVs were electroformed at 10Hz and 1V for 2 hours. After electroformation, GUVs were collected and diluted into an 10mM HEPES pH 7.4, 100 mM NaCl buffer containing 100 nM EqT-Alexa647. GUVs were imaged using a Nikon A1RHD25 confocal microscope equipped with a 40x CFI Plan Apo Lambda S Sil objective.

### FACS-based assay for lipid recycling, retrograde, and lysosomal trafficking

FACS-based assays were conducted as described in Schmieder et al (Schmieder et al., 2022). In brief, cells were plated in 24 well plates to 70% confluency, washed, and equilibrated with serum-free DMEM (without phenol red) for 5min at 37°C. Fluorescent GM1 C18:0 and GM1 C18:1^Δ9^ fusion species were diluted to 0.25 µM or 0.1 µM concentrations of the respective peptide-labeled GM1 species in serum-free DMEM (without phenol red) with molar ratio of dfBSA (1:0.8 lipid:dfBSA). Lipids were loaded for 10 min at 37°C and washed in serum-free DMEM (without phenol red). To measure the amount of incorporated (‘load’) GM1, cells were trypsinized for 5 min at 37°C, then chilled to 4°C and stained with streptavidin-AlexaFluor™ (1:2500, 2 mg/ml) in PBS containing 5% BSA for 15min at 4°C. Cells were washed twice with PBS containing 5% FBS and analyzed using a FACS Canto II (BD, Biosciences, NJ). To obtain recycling or ‘chase’ samples, the GM1 species were washed off in 1x DMEM without phenol red and cells were subsequently incubated for 30 min, 1.5 h, and 3 h in DMEM containing 5% FBS in the presence of 4kDa Dextran-CascadeBlue (500 μg/ml). Cells were trypsinized and stained with streptavidin-AlexaFluor™ as described above.

For lysosomal trafficking assay, cells were chilled to 4°C and stained using 5 nM CTxB-pHrodo™ in PBS containing 5% BSA. Unbound CTxB was washed off and cells were incubated for 1.5 h at 37°C in DMEM containing 5% FBS. Total cell surface GM1 was determined using 5 nM fluorescently labeled CTxB and used to normalize CTxB-pHrodo™ values across the different A431 KO cell lines.

Three biological replicates per assay were performed and 10,000-20,000 single cells were recorded per treatment for all FACS-based assays.

### Lysosomal trafficking of LDL and TFN receptor

A431 CRISPR-Cas9, Cav1, or Cavin-1 KO cell lines were grown to 70% confluency in 24 well plates. To determine the lysosomal trafficking of different membrane protein receptors, cells were washed twice with 1x PBS and chilled to 4°C. Cells were then loaded with respective TFN and LDL at (1 and 5 μg/ml), as either pH sensitive (phrodo™) or pH resistant (AlexaFluor™/Bodipy for LDL). Plasma membrane receptor amount ‘Load’ was determined by incubating pre-chilled cells for 10min at 4°C with pH-resistant ligand. Cells were subsequently washed in PBS and detached using Cell Dissociation Buffer (Gibco). Lysosomal fraction was determined by loading cells with pH-sensitive ligand as above; cells were moved to 37°C for 1.5h in the presence of 4kDa dextran-CascadeBlue to normalize endocytosis rates. Cells were trypsinized and subsequently analyzed by FACS.

### Lipid loading, imaging, and quantification for Airyscan fluorescence microscopy

A431 CRISPR-Cas9, Cav1, or Cavin-1 KO cell lines were plated on glass bottom petri dishes (MatTek, Ashland, MA). Fluorescent GM1 variants were added as described for the FACS-based assays and their subcellular localization was determined after 15 min or 30 min at 37°C in the presence of fluorescent transferrin (1 μg/ml) and LysoTracker™ (2 nM) or 4kDa dextran CascadeBlue (1 mg/ml) to demarcate the respective organelles. Sorting endosomes were defined as vesicles containing transferrin and dextran. Airyscan imaging was performed using a Zeiss LSM 880 microscope with an Airyscan detector and Plan-Apochromat 63x oil immersion objective with (NA = 1.4). Airyscan processing was performed using the Zeiss ZenBlack software. All quantifications were performed using Fiji software (ImageJ, National Institutes of Health, Bethesda, MD, USA).

GM1 in recycling endosomal tubules: A region of interest (ROI) was manually selected, with each containing approximately 3 - 6 triple-positive vesicles containing dextran, transferrin, and GM1. This ROI was thresholded using (Otsu) to separate the background and binarized. GM1 presence/absence (1/0) within transferrin-positive tubules was recorded and % for the ROI was calculated.

Quantification of membrane fluidity in endosomal vesicles and tubules: A431 CRISPR-Cas9, Cav1, or Cavin-1 KO cell lines were plated on glass bottom petri dishes (MatTek, Ashland, MA). Di-4-ANEPPDHQ and Laurdan and fluorescent transferrin (1 μg/ml) were added for 15 min at 37 °C. Sorting endosomes were determined using transferrin and Di-4-ANEPPDHQ or C-Laurdan ratios were measured for each according to (Owen et al., 2011; Sezgin et al., 2012; Steinkuhler et al., 2019). The di-4-ANEPPDHQ generalized polarization spectra (GP) was calculated: 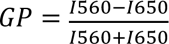

where *I*_560_ and *I*_650_ are the fluorescence intensity emitted at 560 nm and 650 nm, respectively. For Laurdan: GP = (I_440_ − I_490_)/(I_440_ + I_490_) where I_440_ and I_490_ are the fluorescence intensities at 440 nm and 490 nm, respectively.

### Cholesterol staining using filipin III

For immunofluorescence microscopy cells were plated on 35mm glass bottom dishes (Mattek) to 70% confluency. Filipin staining was performed by fixing treated cells in 4 % paraformaldehyde for 1 h and subsequent staining with [0.05 mg/ml] filipin III for 1 h to determine plasma membrane cholesterol contents. Plasma membrane fluorescence intensity was measured using the Fiji (ImageJ, National Institutes of Health, Bethesda, MD, USA) segmented line tool to outline plasma membrane, and mean grey values were obtained.

### GPMV generation and C- Laurdan staining

A431 CRISPR-Cas9, CAV1, or Cavin-1 KO cell lines were seeded two days prior to assay on 12 well plates to reach 70% confluency. Cells were equilibrated by washing twice in GPMV buffer (50 mM Tris, pH = 8, 150 mM NaCl, 50 mM CaCl_2_). Vesiculation was achieved according to (Sezgin et al., 2012) by using 2 mM DTT and 0.07 % PFA in GPMV buffer and cells were incubated for 1 h at 37°C. Supernatant containing GPMV vesicles were harvested and sedimented 5 min/ 500 g and subsequently stained using [0.5 μM] C-laurdan for 3 h at RT. Dye was washed off twice and vesicles were chilled to 10 °C before imaging. Imaging was achieved by using spectral imaging with a confocal microscope Zeiss LSM 880 using Plan-Apochromat 63x oil immersion objective (NA 1.4). C-laurdan was excited at 405 nm and the emission was collected with a multichannel spectral detector (PMT) in the range 430 - 450 nm and 480 - 500 nm simultaneously. The polarity index, generalized polarization (GP) was calculated as described above.

### Lipid extraction and Lipidomic profiling of A431 Cas9 and Cav1-cell lines

Lipidomic profiling was performed as described in Schmieder et al (Schmieder et al., 2022). In brief, 500,000 cells per biological replicate were used. Cells were flash-frozen in liquid nitrogen and subsequently lyophilized. Lipid extraction and lipidomic profiling were performed according to Liaw *et. al*. and Kumar *et. al*. (Gajenthra Kumar et al., 2018; Liaw et al., 2016). In brief, the lyophilized pellet was homogenized in 1 ml ddH_2_O, subsequently, 2.2 ml methanol and 0.9 ml dichloromethane was added. Suspension was incubated o/N at −20C before 20 μl Avanti Splash mix was added, mixture was vortexed and incubated for 10 min at RT. Then 1 ml ddH_2_O, 0.9 ml dichloromethane, and 0.5 ml methanol were added. Solution was centrifuged at 1200 rpm for 10 min and lower phase collected into fresh glass tube. 2 ml dichloromethane was added to the aqueous phase to extract any remaining lipids, mixture was centrifuged as above and the organic phase was pooled with the first one. Samples were dried down in SpeedVac.

Unbiased MS/MSALL shotgun lipidomic profiling was performed on a Sciex 5600 TripleTOF mass spectrometer.

### Phylogenetic analysis of Caveolin-1 cysteines

The distribution of Cys residues within the spoke region was performed using recently reported phylogenetic analysis of 74 caveolins (Han et al., 2025). Probability analysis for the enrichment of cysteine residues across the caveolin paralogs was performed using custom-based code. Probability for the frequency of a cysteine across ten amino acids was measured and plotted across the different paralogs. To infer the evolutionary relationships among chordate caveolin homologs, we employed a maximum likelihood (ML) approach using IQ-TREE (v2.1.4-beta COVID-edition). Protein sequences were aligned and filtered using GUIDANCE2 (v2.0) within a Docker container (kingcohn1/guidance2) with MAFFT as the alignment program, 100 bootstrap iterations, and default confidence thresholds to remove poorly aligned regions. The input sequences were pre-processed to remove non-standard characters and empty lines, ensuring compatibility with GUIDANCE2. The filtered alignment (MSA.MAFFT.Without_low_SP_Col.With_Names) was then analyzed with IQ-TREE in a Google Colab environment equipped with a Conda-based bioconda channel. IQ-TREE was run using the -m MFP option for automatic model selection via ModelFinder, with branch support assessed using 1000 ultrafast bootstrap replicates (-bb 1000) and 1000 SH-aLRT tests (-alrt 1000). Analyses were performed with a single thread (-nt 1) and a fixed random seed (-seed 42) for reproducibility. The resulting phylogenetic tree was visualized using the Interactive Tree of Life (iTOL) platform.

### AlphaFold v3.0 predictions

AlphaFold v3.0 predictions were performed using the AlphaFold Server (https://alphafoldserver.com/) (Abramson et al., 2024).

### Software and Statistical analysis

Imaging data was analyzed using either Zeiss ZENblack or Fiji software. Mean grey fluorescent values of microscopy experiments or mean fluorescence data obtained from FACS experiments (FlowJo, LLC) were transferred to GraphPad Prism software (San Diego, CA) for graphing and statistical analysis. Statistically significant difference between treatments was tested by One-way ANOVA with Tukey’s multiple comparisons; alternatively, an unpaired t-test was applied when indicated with n.s. p > 0.05, *p < 0.05, **p < 0.01 ***p < 0.001 and ****p < 0.0001.

---

**Suppl. Figure 1.**
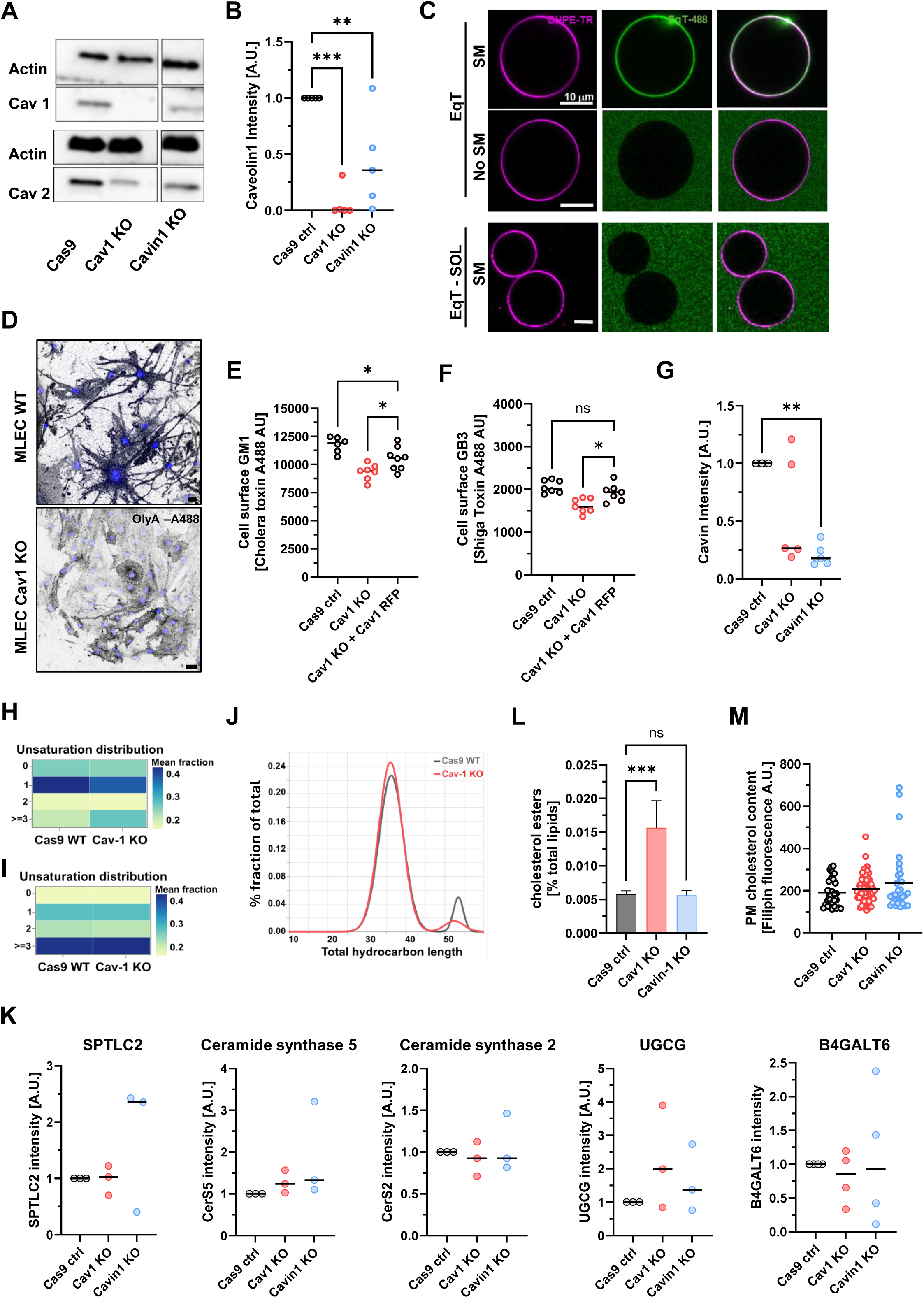
Cell surface depletion of complex sphingolipids in the absence of Cav1. **(A)** Validation of A431 Cav1 and Cavin-1 KOs. Cav1 is not detected by Western blot in A431 Cav1 KO and slightly reduced in A431 Cavin-1 KO. A431 Cav1 and Cavin-1 KO both show reduction in Cav2. **(B)** Western blot quantification of (A), normalized to actin. **(C)** EqT-A488 binding to GUVs containing or not SM. GUVs are labeled with membrane marker DHPE-TR. Non-binding EqT (EqT-SOL) does not bind to GUVs containing SM. Scale bar = 10 µm. **(D)** Cell surface binding of SM binding OlyA-A647 to MLEC WT and Cav1 KO. DAPI in blue. Scale bar = 20 µm. **(E)** FACS analysis of cell surface GM1 by CTxB- A488 in A431 Cas9 WT, Cav1 KO or Cav1 KO cells overexpressing Cav1-RFP. N = 5 biological replicates, dot represents 20.000 cells. **(F)** FACS analysis of cell surface GB3 by STxB-A488 in A431 Cas9 WT, Cav1 KO or Cav1 KO cells overexpressing Cav1-RFP. N = 7 biological replicates, dot represents 20.000 cells. **(G)** Western blot quantification of Cavin-1 expression in Cav1 and Cavin-1 KO, normalized to actin. **(H)** Lipidomics analysis of A431 Cas9 WT or Cav1 KO cells, n = 3 biological replicates. Heatmap of mean fraction displaying degree of unsaturation in total SM between A431 Cas9 WT and A431 Cav1 KO. Unsaturation of ≥3 statistically different. **(I)** Heatmap displaying mean fraction for the degree of unsaturation across total lipids is not statistically different. **(J)** Chain length distribution profile of the total lipids in A431 Cas9 WT (grey) or A431 Cav1 KO (red). Acyl chain carbon number 30 to 36 and 46-58 are significantly different. **(K)** Western blot analysis for genes involved in the GSL and SL pathways between A431 Cas9 WT, Cav1 KO and Cavin-1 KO. **(L)** Total amount of cholesterol esters is increased in A431 Cav1 KO compared to WT or Cavin-1 KO. **(M)** Plasma membrane cholesterol content measured by filipin staining for A431 Cas9 WT, Cav1 KO and Cavin-1 KO. For all experiments: n.s. p > 0.05, *p < 0.05, **p < 0.01 ***p < 0.001 and ****p < 0.0001.

**Suppl. Figure 2.**
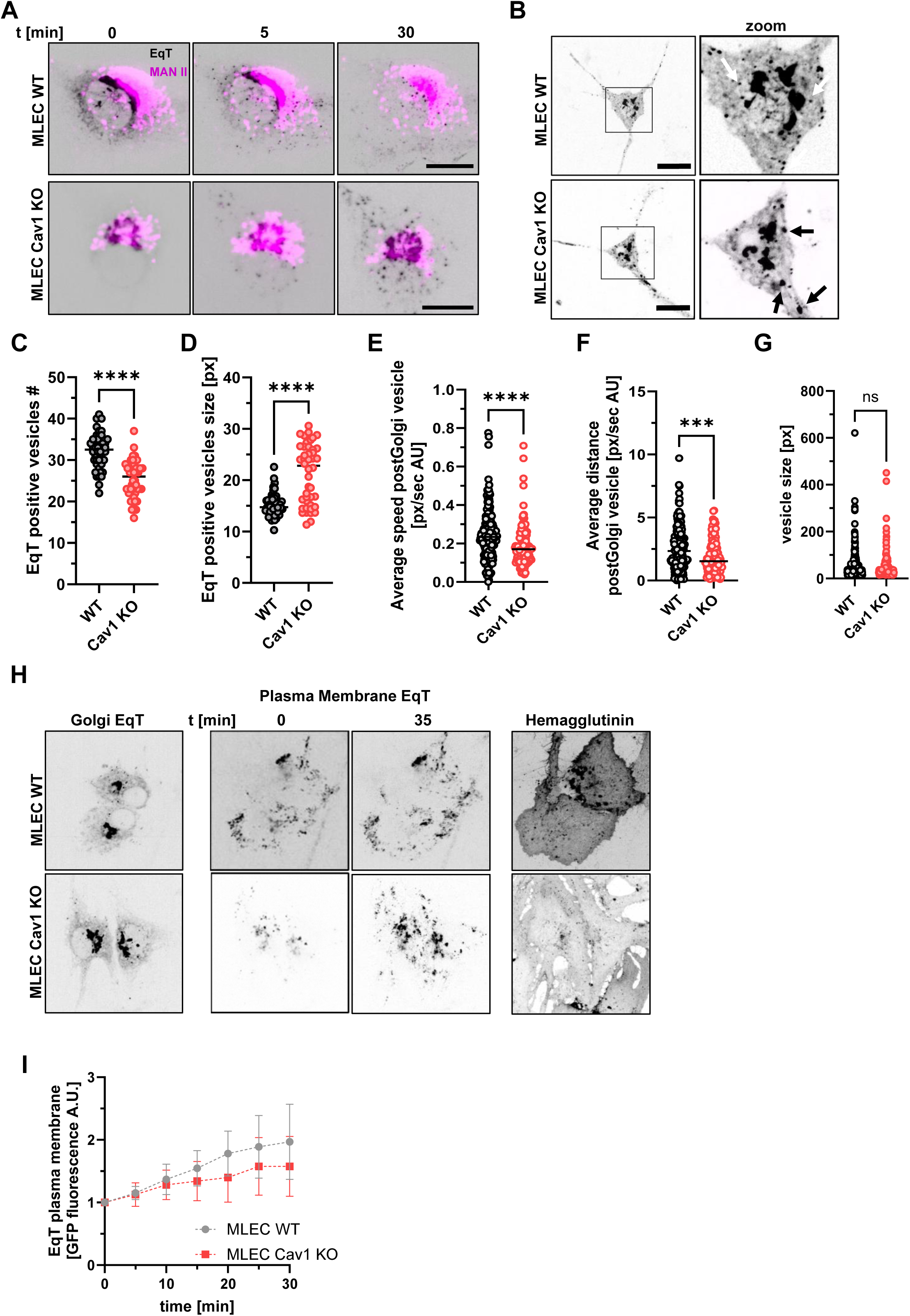
MLEC Cav1 KO cells display aberrant Golgi and secretory post-Golgi morphologies and moderately slower secretory kinetics for SM. **(A)** RUSH experiment Golgi mask using the cis Golgi marker αManII (magenta). Maximal co-localization between EqT-RUSH and αManII defines time point 0. Scale bar = 10 µm. **(B)** Post-Golgi EqT vesicles in MLEC WT and Cav1 KO cells. Scale bar = 20 µm **(C)** Quantification of number of EqT positive post-Golgi vesicles in MLEC WT and Cav1 KO cells. N= 2 biological replicates. **(D)** Quantification of size of EqT positive post-Golgi vesicles in MLEC WT and Cav1 KO cells. N= 2 biological replicates. **(E)** Quantification of post-Golgi EqT vesicles, measuring vesicle speed in MLEC WT and Cav1 KO cells. N = 2 biological replicates. **(F)** Quantification of post-Golgi EqT vesicle dynamics, tracking total distances of individual vesicles in MLEC WT and Cav1 KO cells. N = 2 biological replicates. **(G)** Quantification of post-Golgi EqT-sol vesicles, measuring vesicle speed in MLEC WT and Cav1 KO cells. N = 2 biological replicates. **(H)** Time course of PM and intracellular EqT-RUSH distribution in MLEC WT and Cav1 KO cells. **(I)** Quantification of time course for PM staining of EqT in MLEC WT and Cav1 KO cells in (H). N = 3 biological replicates. For all experiments: n.s. p > 0.05, *p < 0.05, **p < 0.01 ***p < 0.001 and ****p < 0.0001.

**Suppl. Figure 3.**
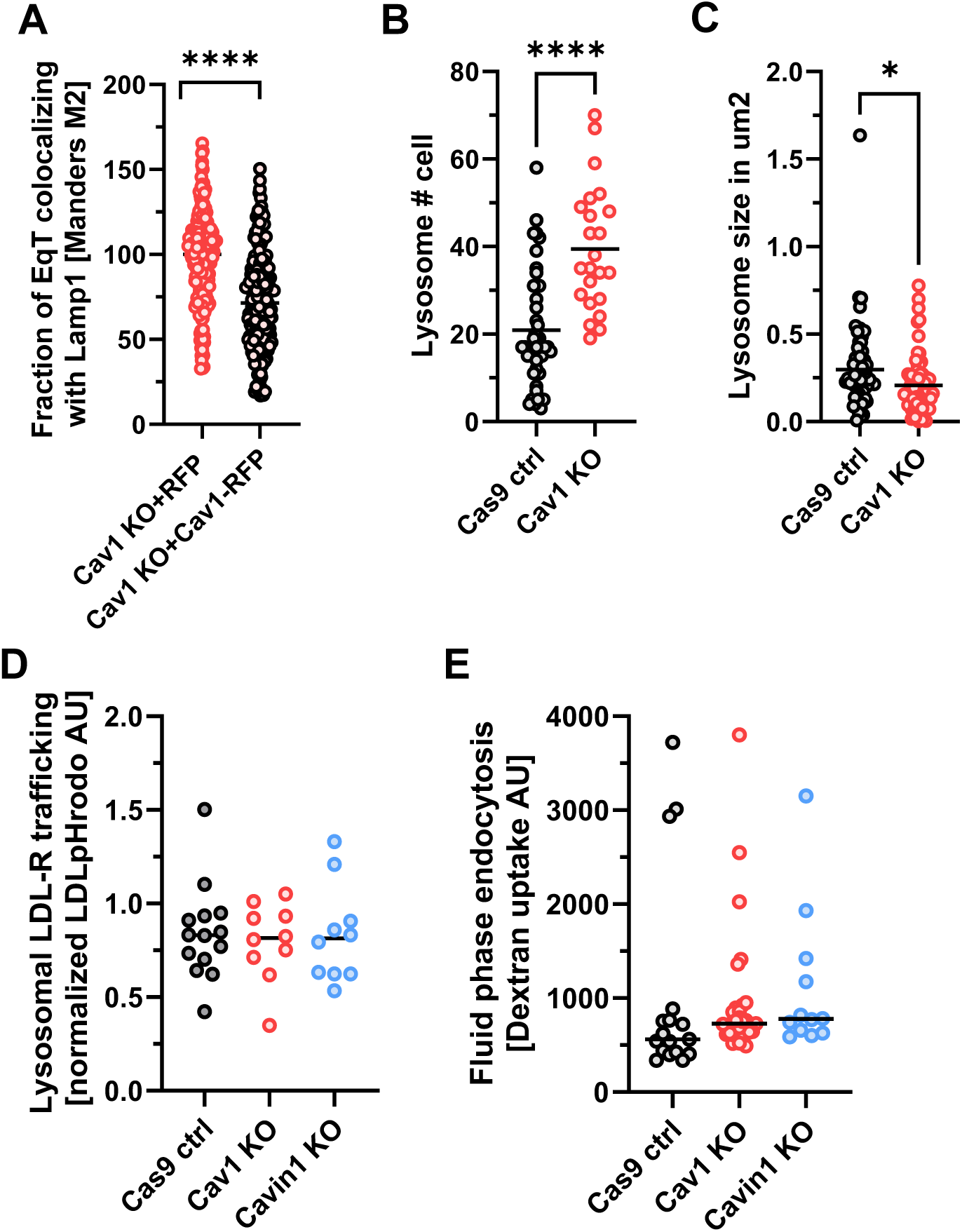
Redistribution of SLs to lysosomes in Cav1 KO cells. **(A)** Overexpression of Cav1- RFP but not RFP alone in MLEC Cav1 KO leads to redistribution of intracellular SM measured by co- localization of SM EqT-A488 staining with lysosomal marker Lamp-1. Data normalized for WT to 100. Mander’s M2 quantification of co-localization between EqT-A488 and Lamp-1 for n = 3 biological replicates. **(B)** Quantification of lysosome number in A431 Cas9 WT and Cav1 KO cells. N ≥4 biological replicates. **(C)** Quantification of lysosome size in A431 Cas9 WT and Cav1 KO cells. N ≥ 4 biological replicates. **(D)** Lysosomal trafficking of the lysosomal cargo LDL-R by LDL-pHrodo measured by FACS is not impaired in A431 Cav1 KO or Cavin-1 KO compared to A431 Cas9 WT. N ≥ 10 biological replicates, dots represent 20.000 cells. **(E)** Fluid phase uptake of Dextran-Pacific Blue measured by FACS is not different in A431 Cas9 WT, Cav1 KO and Cavin-1 KO. N ≥ 10 biological replicates, dots represent 20.000 cells. For all experiments: n.s. p > 0.05, *p < 0.05, **p < 0.01 ***p < 0.001 and ****p < 0.0001.

**Suppl. Figure 4.**
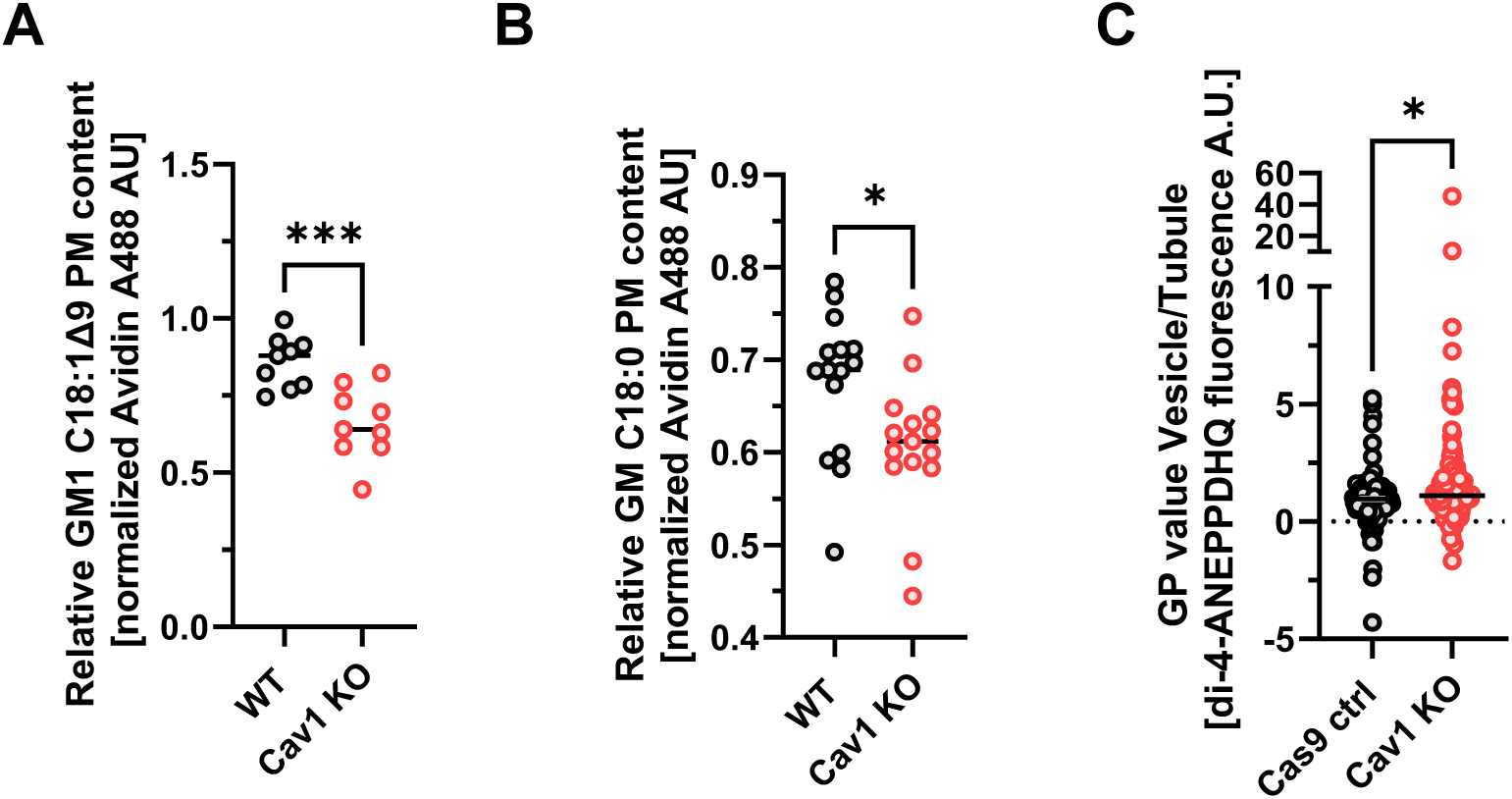
Sphingolipid plasma membrane recycling is impaired in Cav1 KO. **(A)** Quantification of plasma membrane recycling of GM1 C18:1^Δ9^ in MLEC WT and Cav1 KO after 1.5h. N= 3 biological replicates, dots represent 20.000 cells. **(B)** Quantification of plasma membrane recycling for GM1 C18:0 in MLEC WT and Cav1 KO after 1.5h. N= 3 biological replicates, dots represent 20.000 cells. **(C)** Quantification of the membrane order (GP value) as ratio between recycling endosomal vesicle and tubules measured by the polarity sensitive dye di-4-ANEPPDHQ in TFN positive recycling vesicles and tubules for A431 Cas9 WT and Cav1 KO is increased in A431 Cav1 KO. N ≥ 4 biological replicates. For all experiments: n.s. p > 0.05, *p < 0.05, **p < 0.01 ***p < 0.001 and ****p < 0.0001.

**Suppl. Figure 5.**
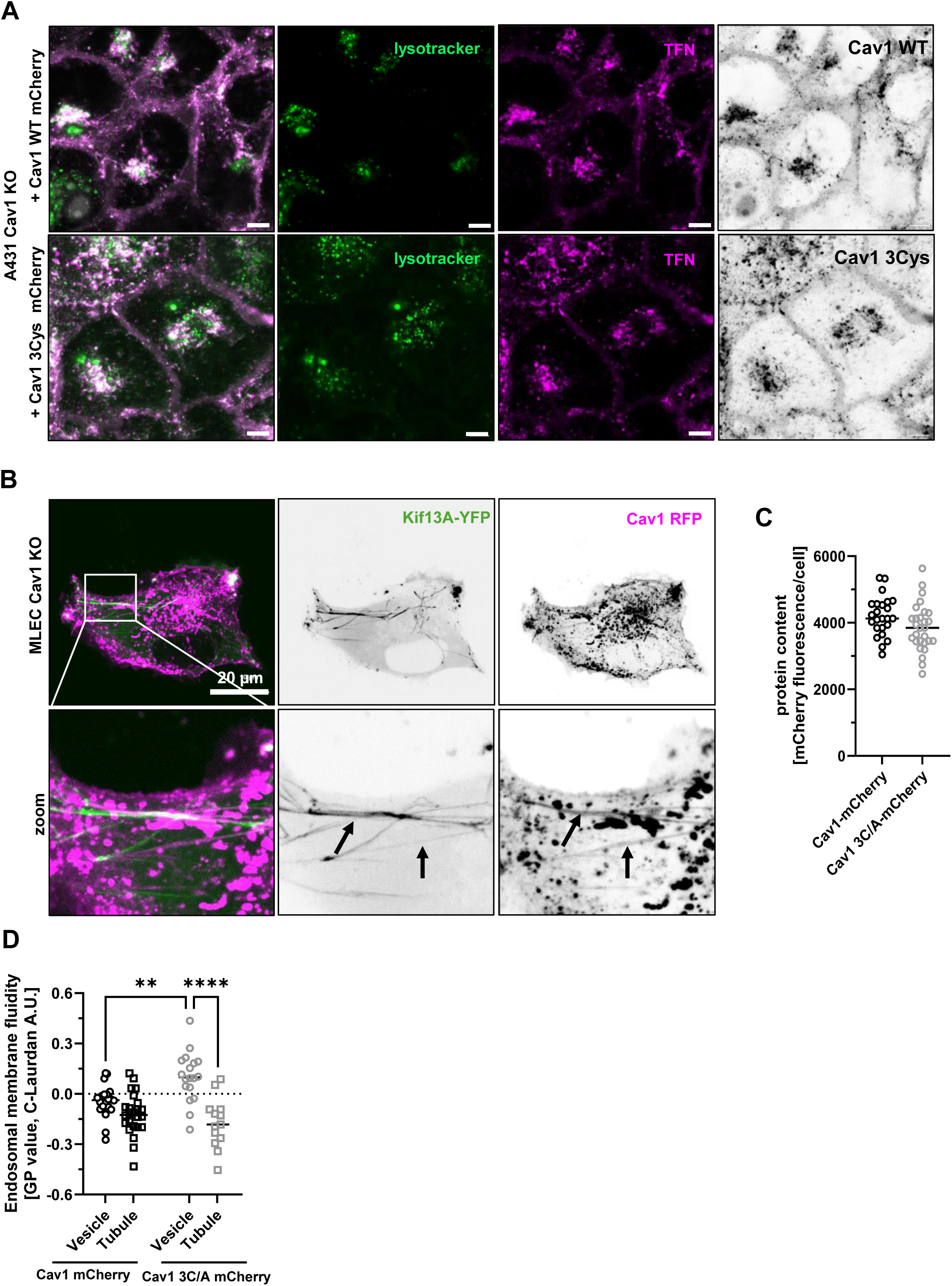
Cav1 palmitoylations are required for endosomal SL sorting. **(A)** Subcellular localization of Cav1 WT (top panels) and Cav1-3Cys mutant protein (bottom panels) is the same in A431 Cav1 KO cells overexpressing either Cav1 WT mCherry or Cav1-3Cys mutant mCherry protein. Scale bar = 20 µm. **(B)** MLEC Cav1 KO overexpressing dominant positive Kif13A-YFP and Cav1-RFP displays co- localization of Cav1-RFP to Kif13A positive recycling tubules. Scale bar = 20 µm. **(C)** Quantification of fluorescence per cell for A431 Cav1 KO cell overexpressing either Cav1-mCherry or Cav1-3C/A mutant-mCherry. Dots represent 20.000 cells. N= >5 biological replicates. **(D)** Quantification of membrane order (GP value) measured by the polarity sensitive dye Laurdan in TFN positive recycling vesicles and tubules for A431 Cav1 KO overexpressing either Cav1 WT mCherry or Cav1-3C/A mutant mCherry. Cav1-3C/A mutant protein cannot rescue the decreased order in recycling tubules of A431 Cav1 KO. N ≥ 4 biological replicates. For all experiments: n.s. p > 0.05, *p < 0.05, **p < 0.01 ***p < 0.001 and ****p < 0.0001.

**Suppl. Figure 6.**
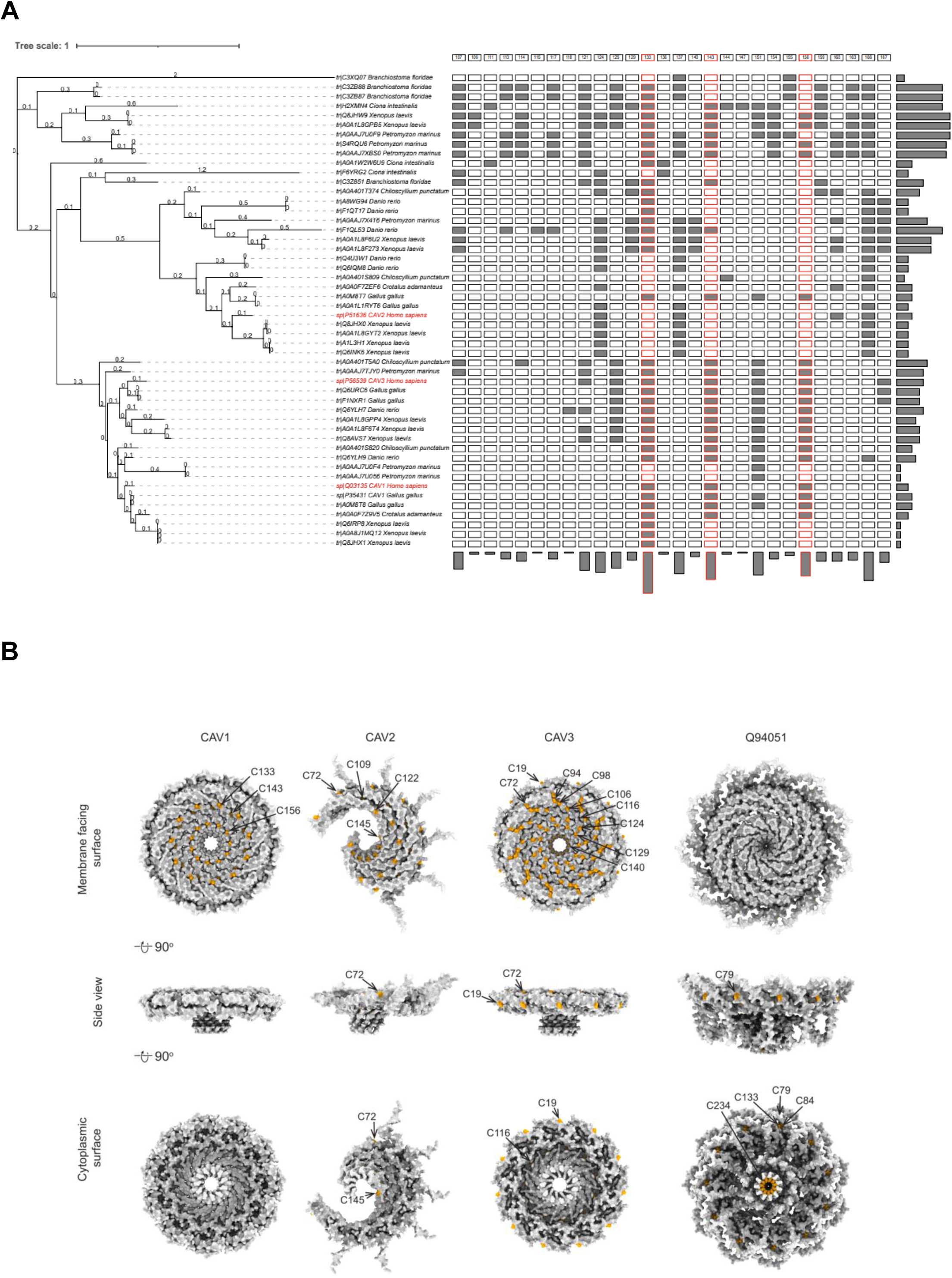
Evolutionary analysis of human Cav1 palmitoylation sites. **(A)** Conservation of C133, C143, and C156 in human Cav1 (red box) across the chordate Cav homologs. Human Cav1, 2 and 3 in (red). **(B)** Alphafold rendering of human Cav1, Cav2 and Cav3, as well as CeCav displaying cysteine residues for palmitoylation (yellow) in the spoke region. Displayed are proteins from membrane facing surface (top row), side view (middle row) and from the cytosolic side (bottom row).

